# Existence and functions of hypothalamic kisspeptin neuropeptide signaling system in a non-chordate deuterostome species

**DOI:** 10.1101/851261

**Authors:** Tianming Wang, Zheng Cao, Zhangfei Shen, Jingwen Yang, Xu Chen, Zhen Yang, Ke Xu, Xiaowei Xiang, Qiuhan Yu, Yimin Song, Weiwei Wang, Yanan Tian, Lina Sun, Libin Zhang, Su tGuo, Naiming Zhou

**Author notes:** These authors contributed equally: Tianming Wang, Zheng Cao and Zhangfei Shen. Correspondence and requests for materials should be addressed to N.Z. or to T.W.

## Abstract

The kisspeptin (Kp) system is a central modulator of the hypothalamic-pituitary-gonadal axis in vertebrates. Its existence outside the vertebrate lineage remains largely unknown. Here we report the identification and characterization of Kp system in the sea cucumber *Apostichopus japonicus.* The gene encoding the Kp precursor, generates two mature neuropeptides, AjKiss1a and AjKiss1b. The Kp receptors, AjKissR1 and AjKissR2, are strongly activated by synthetic *A. japonicus* and vertebrate Kps, triggering a rapid intracellular mobilization of Ca^2+^, followed by receptor internalization. AjKissR1 and AjKissR2 share similar intracellular signaling pathways via G_αq_/PLC/PKC/MAPK cascade, when activated by C-terminal decapeptide (AjKiss1b-10). The *A. japonicus* Kp system functions in mutiple tissues which are closely related to reproduction and metabolism. Overall, our findings uncover for the first time, to our knowledge, the existence and function of the Kp system in a non-chordate species and provide new evidence to support the ancient origin of the hypothalamic neurosecretory system.

## Introduction

Nervous systems, from simple nerve nets in primitive species to complex architectures in vertebrates, process sensory stimuli and enable animals to generate body-wide responses [1]. Neurosecretory centers, one of the major output systems in the animal brain, secrete neuropeptides and nonpeptidergic neuromodulators to regulate developmental and physiological processes [2]. Understanding the evolutionary origin of these centers is an area of active investigation, mostly because of their importance in a range of physical phenomena such as growth, metabolism, or reproduction [3, 4].

The hypothalamus constitutes the major part of the ventral diencephalon in vertebrates and acts as a neurosecretory brain center, controlling the secretion of various neuropeptides (hypothalamic neuropeptides) [5, 6]. Outside vertebrates, similar neurosecretory systems have been seen in multiple protostomian species including crustaceans, spiders, and molluscs [7]. Specific to echinoderms, which occupy an intermediate phylogenetic position as a deuterostomian invertebrate species with respect to vertebrates and protostomes, increasing evidence, collected from *in silico* identification of hypothalamic neuropeptides and functional characterization of vasopressin/ocytocin (VP/OT)-type signaling system, suggests the existence of a conserved neurosecretory system [4, 8].

The hypothalamic neuropeptide kisspeptins (Kps), encoded by the *Kiss1* gene and most notably expressed in the hypothalamus, share a common Arg-Phe-amide motif at their C-termini and belong to the RFamide peptide family [9, 10]. Exogenous administration of Kps triggers an increase in circulating levels of gonadotropin-releasing hormone and gonadotropin in humans, mice, and dogs [11-14]. Accumulating evidence suggests that the Kp system functions as a central modulator of the hypothalamic-pituitary-gonadal (HPG) axis to regulate mammalian puberty and reproduction through a specific receptor, GPR54 (also known as AXOR12 or hOT7T175), which is currently referred to as the Kp receptor (KpR) [15-17]. Following the discovery of Kps and KpRs in mammals, a number of Kp and KpR paralogous genes have been revealed in other vertebrates [18], and a couple of functional Kp/KpR have also been demonstrated in amphioxus [19]. Moreover, Kp-type peptides and their corresponding receptors, in echinoderms, have been annotated *in silico*, based on the analysis of genome and transcriptome sequence data [20-25]. However, to our knowledge, neither the Kp-type peptides nor the corresponding receptors have been experimentally identified and functionally characterized in non-chordate invertebrates. This raises an important question: does the Kp/KpR signaling system have an ancient evolutionary origin or did it evolve *de novo* in the chordate/vertebrate lineages?

Here, we addressed this question by searching for Kp/KpR genes in a non-chordate species, the sea cucumber *Apostichopus japonicas*. It is one of the most studied echinoderms and is widely distributed in temperate habitats in the western North Pacific Ocean, being cultivated commercially on a large scale in China [26]. We uncovered *Kiss-like* and *KissR-like* genes by mining published *A. japonicus* data [25, 27], using a bioinformatics approach. Their signaling properties were characterized using an *in vitro* culture system. Through the evaluation of Ca^2+^ mobilization and other intracellular signals, we found that *A. japonicus* Kps dramatically activated two Kp receptors (AjKissR1 and AjKissR2), via a GPCR-mediated G_αq_/PLC/PKC/MAPK signaling pathway, that have functions corresponding to those of the vertebrate Kp system. Finally, we revealed the physiological activities of this signaling system both *in vivo* and *ex vivo*, and we demonstrated the involvement of the Kp system in reproductive and metabolic regulation in *A. japonicus*. Collectively, our findings indicate the existence of a Kp/KpR signaling system in non-chordate deuterostome invertebrates and provide new evidence to support the ancient evolutionary origin of the hypothalamic neurosecretory system [3].

## Results

### *In silico* identification of Kps and Kp receptors

Invertebrate Kp receptors have rarely been reported. Putative Kp precursors have been predicted in echinoderms, including starfish (*Asterias rubens*), sea urchin (*Strongylocentrotus purpuratus*), and sea cucumbers (*Holothuria scabra*, *Holothuria glaberrima,* and most recently in *A. japonicus*) [22-25]. Based on these sequences, the putative *A. japonicus* Kp precursor gene was identified *in silico* from transcriptome data and cloned from ovarian tissue samples by reverse transcription polymerase chain reaction (RT-PCR). The full-length cDNA (GenBank accession number MH635262) was 2,481 bp long and contained a 543 bp ORF, encoding a 180 amino acid peptide precursor with one predicted signal peptide region and four cleavage sites (Fig. 1A and Figure 1–figure supplement 1). Two mature peptides with amide donors for C-terminal amidation, 32 amino acid Kp-like peptide with a disulfide-bond (AjKiss1a) and 18 amino acid Kp-like peptide (AjKiss1b), were predicted (Supplementary Table 1). Alignment of multiple sequences revealed a high similarity between AjKiss1a/b and predicted echinoderm Kps but low identity between AjKiss1a/b and vertebrate Kiss1/2 (Fig. 1B). A maximum likelihood tree of Kp precursors, as well as PrRP, 26RFa/QRFP, GnIH, and NPFF from outgroups [28], was constructed for phylogenetic analysis. It showed that the *A. japonicus* Kp precursor, AjKisspeptin, together with kisspeptin-like precursors from the sea cucumbers, *H. scabra* and *H. glaberrima*, were grouped with the vertebrate Kiss1 and Kiss2 subfamilies into the „Kisspeptin‟ group (Fig. 1C).

**Figure 1.**
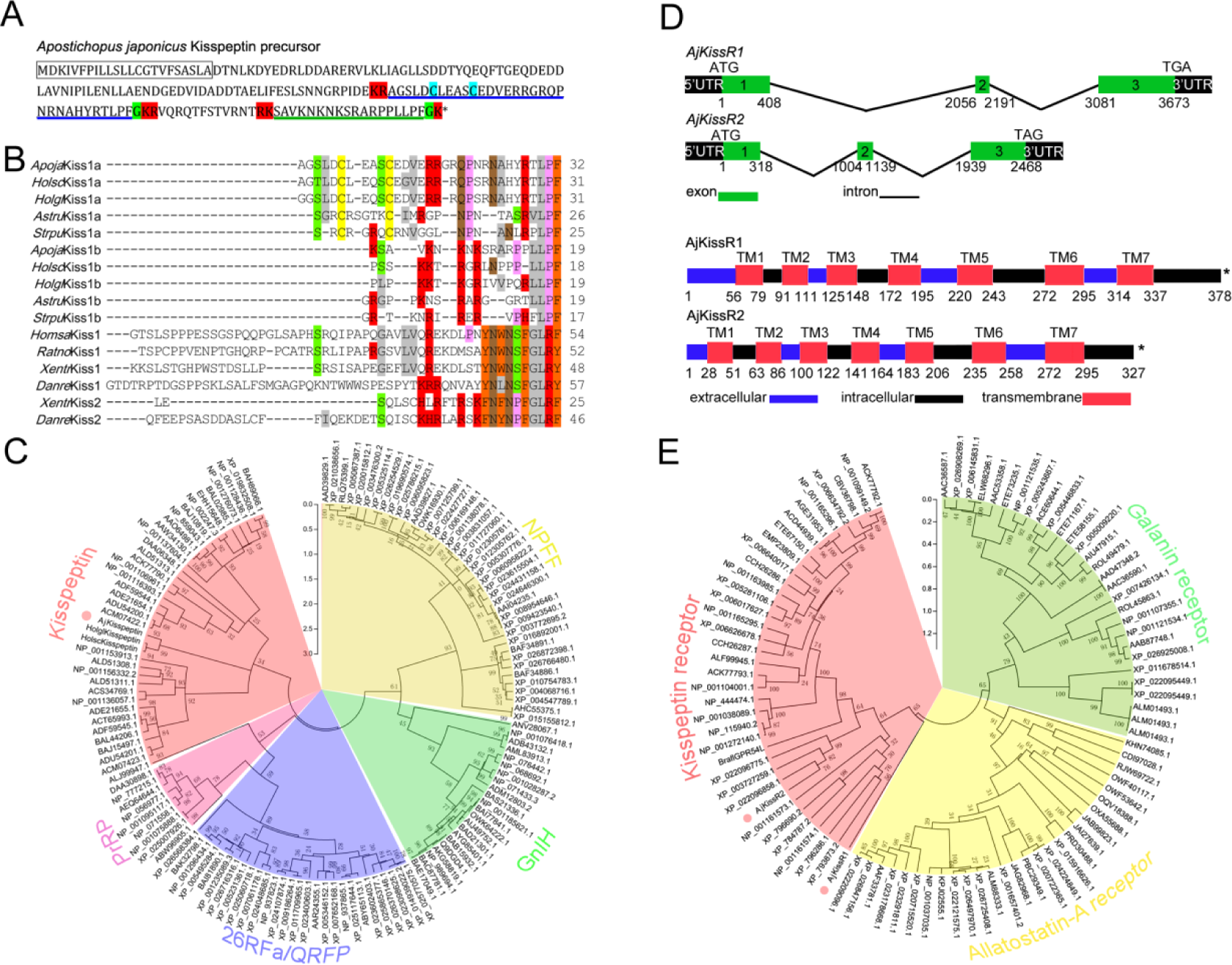
Gene structure, homology, phylogenetic characterization of *Apostichopus japonicus* kisspeptin precursor and kisspeptin receptors. **A.** Deduced amino acid sequence of *A. japonicus* kisspeptin (Kp) precursor. The signal peptide is labeled in box with full lines; the cleavage sites are highlighted in red; glycine residues responsible for C-terminal amidation are highlighted in green; cysteines paired in a disulfide-bonding structure are highlighted in light blue; the predicted mature peptides, AjKiss1a and AjKiss1b, are noted by the blue and green underlines. **B.** Alignment of the predicted echinoderm Kp core sequences and functionally characterized chordate Kps. Sequences of *Holothuria scabra*, *Holothuria glaberrima, Strongylocentrotus purpuratus,* and *Asterias rubens* Kps were predicted by Elphick‟s lab [22, 23]. Vertebrate Kp core sequences were obtained from GenBank with detailed sequences listed in figure 1 raw data set 1. Color align property was generated using Sequence Manipulation Suite online. Percentage of sequences that must agree for identity or similarity coloring was set as 40%. **C.** Phylogenetic tree based on amino acid of kisspeptin precursor and other four different neuropeptide outgroups [28]. The tree was constructed based on maximum likelihood algorithms using MEGA 5.1. The detailed sequences are listed in figure 1 raw data set 2. **D.** DNA and protein structures of AjKissR1/2. *AjKissR1/2* DNA structure is shown with exons numbered in green bands. ATG represents the start methionine codon and TGA/TAG represents the stop codon. Organization of the predicted protein structures is shown. The seven transmembrane domains (TM1– TM7) are marked with red boxes. The N-terminal region and three extracellular (EC) rings are noted with blue boxes, as well as the C-terminal part and three intracellular (IC) rings are indicated with black boxes. Stop codons are represented by an asterisk. Arabic numbers under the band indicate the nucleotide or amino acid sites. **E.** Maximum-likelihood trees of kisspeptin (red), allatostatin-A (yellow) and galanin (green) receptors. The tree was constructed by MEGA 5.1 using allatostatin-A and galanin receptors as outgroups [20]. The detailed sequences are listed in figure 1 raw data set 3. The topological stability of these ML trees was achieved by running 1000 bootstrapping replications. Bootstrap values (%) are indicated by numbers at the nodes.

Several predicted „G-protein coupled receptor 54-like‟ or „kisspeptin receptor-like‟ gene annotations in the hemichordate *Saccoglossus kowalevskii* (two genes), the echinoderm *Acanthaster planci* (two genes), and *S. purpuratus* (seven genes) have been reported [29-31]. Using these predicted genes as reference sequences to search the *A. japonicus* genomic database, three *A. japonicus* Kp receptor-like genes (*AjKissR1, AjKissR2*, and *AjKissRL3*; GenBank accession numbers, MH709114, MH709115, and MG199220, respectively) were identified and cloned from *A. japonicus* ovary by RT-PCR. The open reading frames (ORFs) of both *AjKissR1* and *AjKissR2* (detailed data for *AjKissRL3* have not been presented because it exhibited no interaction with ligands in further experiments) comprised three exons, with deduced amino acid sequences of 378 and 327 residues and contained seven transmembrane domains (Fig. 1D and Figure 1–figure supplement 2). A sequence alignment of AjKissR1 and AjKissR2, with the well characterized chordate GPR54, was performed (Figure 1–figure supplement 3) and a relatively high identity in seven transmembrane region sequences, against 21 vertebrate GPR54 sequences was observed, as shown in Figure 1–figure supplement 4. Maximum likelihood phylogenetic tree analysis, using „Allatostatin-A receptor‟ and „Galanin receptor‟ as outgroups, revealed that AjKissR1 and 2 both clustered in the “Kisspeptin receptor” group. AjKissR2 clustered with the predicted *A. planci* (starfish) Kp receptors (Genbank ID: XP_022096858.1 and XP_022096775.1) and with the *S. purpuratus* (sea urchin) Kp receptor (XP_003727259.1), while AjKissR1 did not group with any known Kp receptors (Fig. 1E).

**Figure 2.**
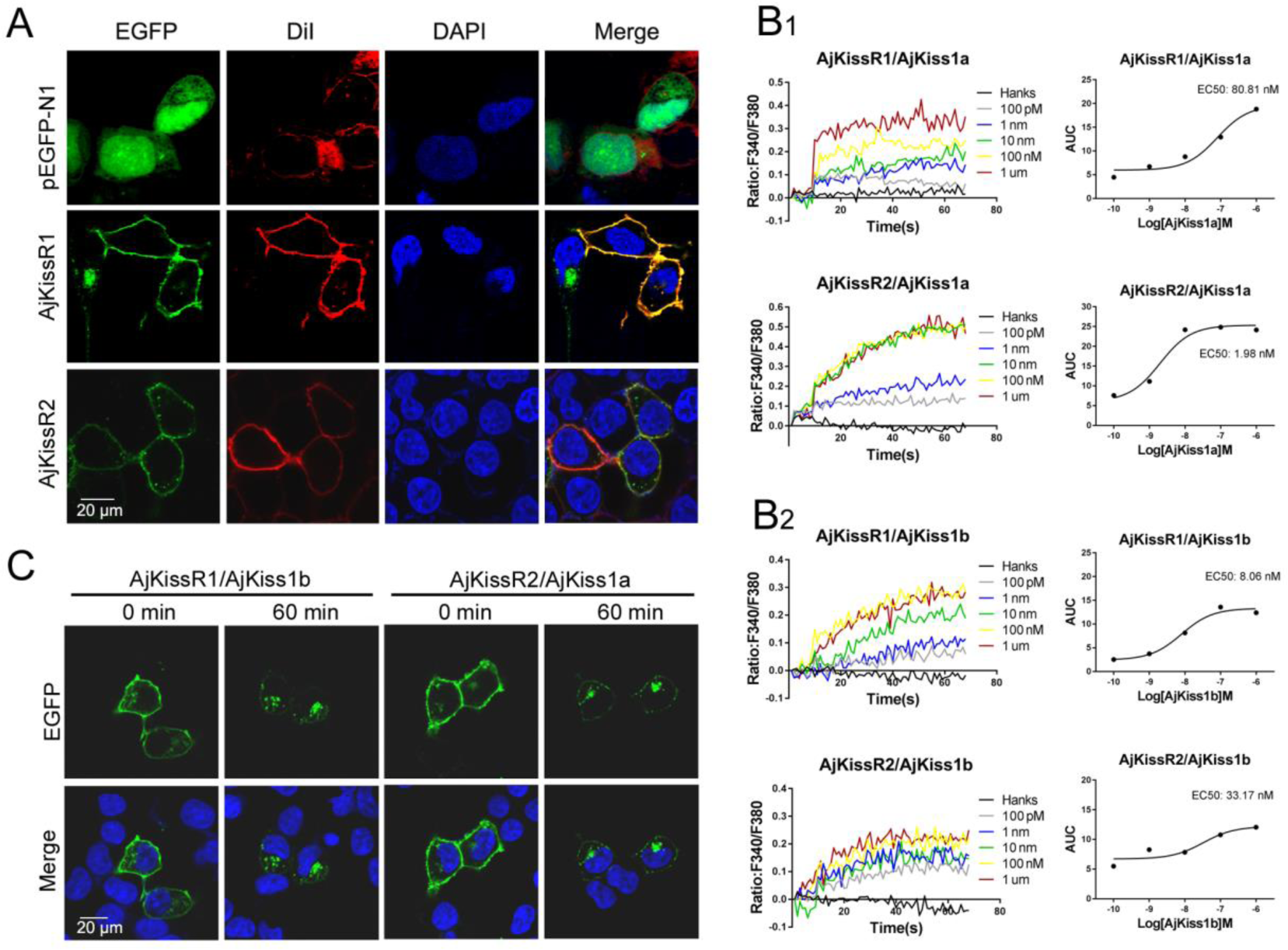
Functional characteristics of *Apostichopus japonicus* kisspeptins (Kps) and receptors. **A.** Transiently expressing AjKissR1-EGFP or AjKissR2-EGFP cells were stained with cell membrane probe (DiI) and cell nucleus probe (DAPI) and detected by confocal microscopy. **B.** Intracellular Ca^2+^ mobilization in flag-AjKissR1 or flag-AjKissR2 expressing HEK293 cells was measured in response to indicated concentrations of AjKiss1a **(B1)** and AjKiss1b **(B2)** using Fura-2/AM, with concentration-dependent course of AjKiss1a or AjKiss1b stimulating Ca^2+^ mobilization in cells. **C.** Internalization of AjKissR1-EGFP or AjKissR2-EGFP initiated by 1.0 μM AjKiss1b in stable AjKissR1-EGFP or 1.0 μM AjKiss1a in stable AjKissR2-EGFP expressing HEK293 cells determined after a 60-min incubation by confocal microscopy.

**Figure 3.**
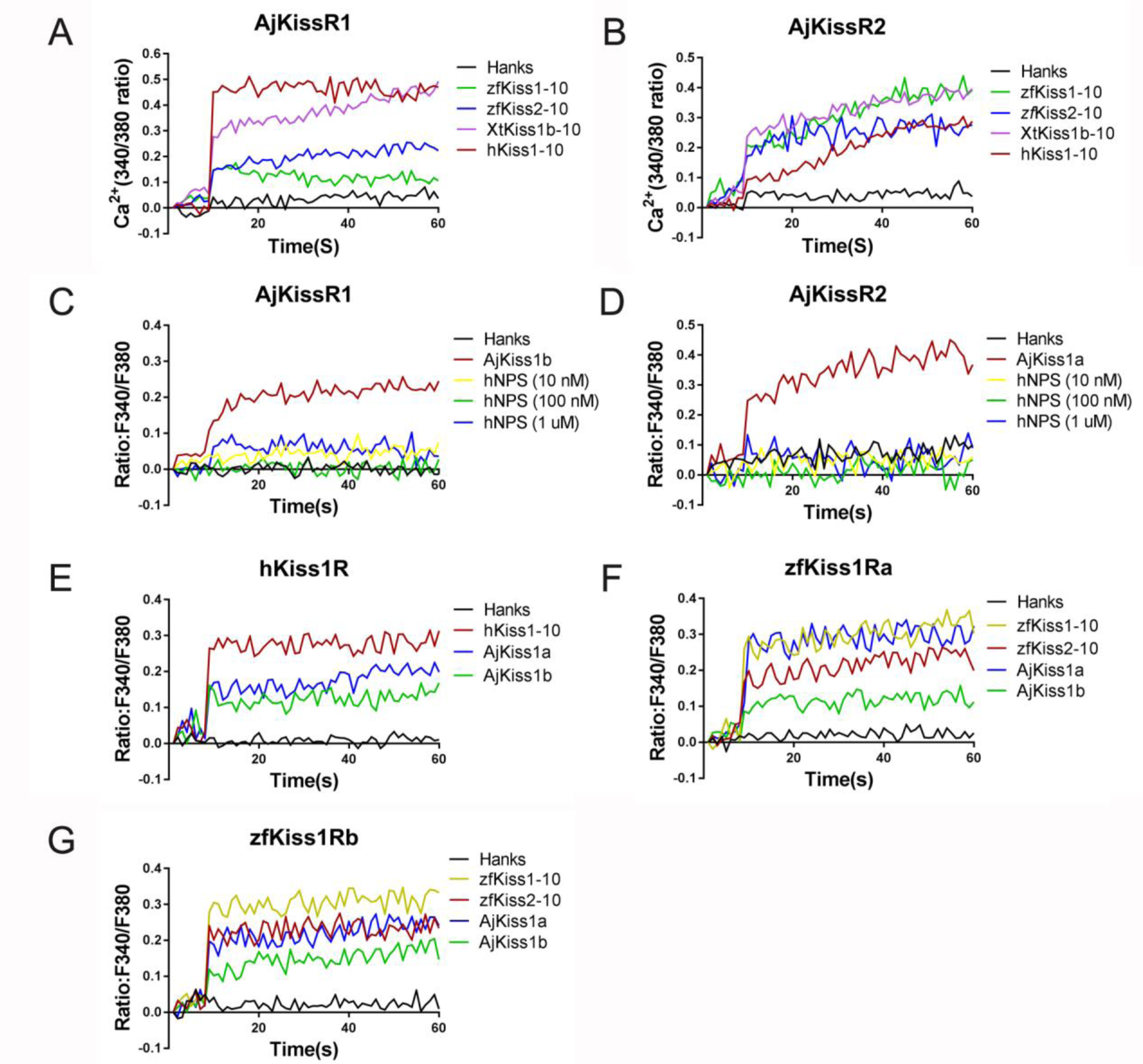
Functional cross-talk between the *A. japonicus* and vertebrate Kisspeptin/Kisspeptin receptor systems. Intracellular Ca^2+^ mobilization in AjKissR1 (**A**) or AjKissR2 (**B**) expressing HEK293 cells was measured in response to 1.0 μM zfKiss1-10, zfKiss2-10, XtKiss1b-10, or hKiss1-10 using Fura-2/AM. No Ca^2+^ mobilization-mediated activity was detected in AjKissR1 (**C**) or AjKissR2 (**D**) expressing HEK293 cells upon administration of indicated concentrations of human neuropeptide S (hNPS). Intracellular Ca^2+^ mobilization in human kisspeptin (Kp) receptor (hKiss1R) expressing HEK293 cells was measured in response to 1.0 μM hKiss1-10, AjKiss1a or AjKiss1b (**E**), as well as in zebrafish Kp receptor (zfKiss1Ra or zfKiss1Rb) expressing cells responding to 1.0 μM zfKiss1-10, AjKiss1a, or AjKiss1b (**F**, **G**).

**Figure 4.**
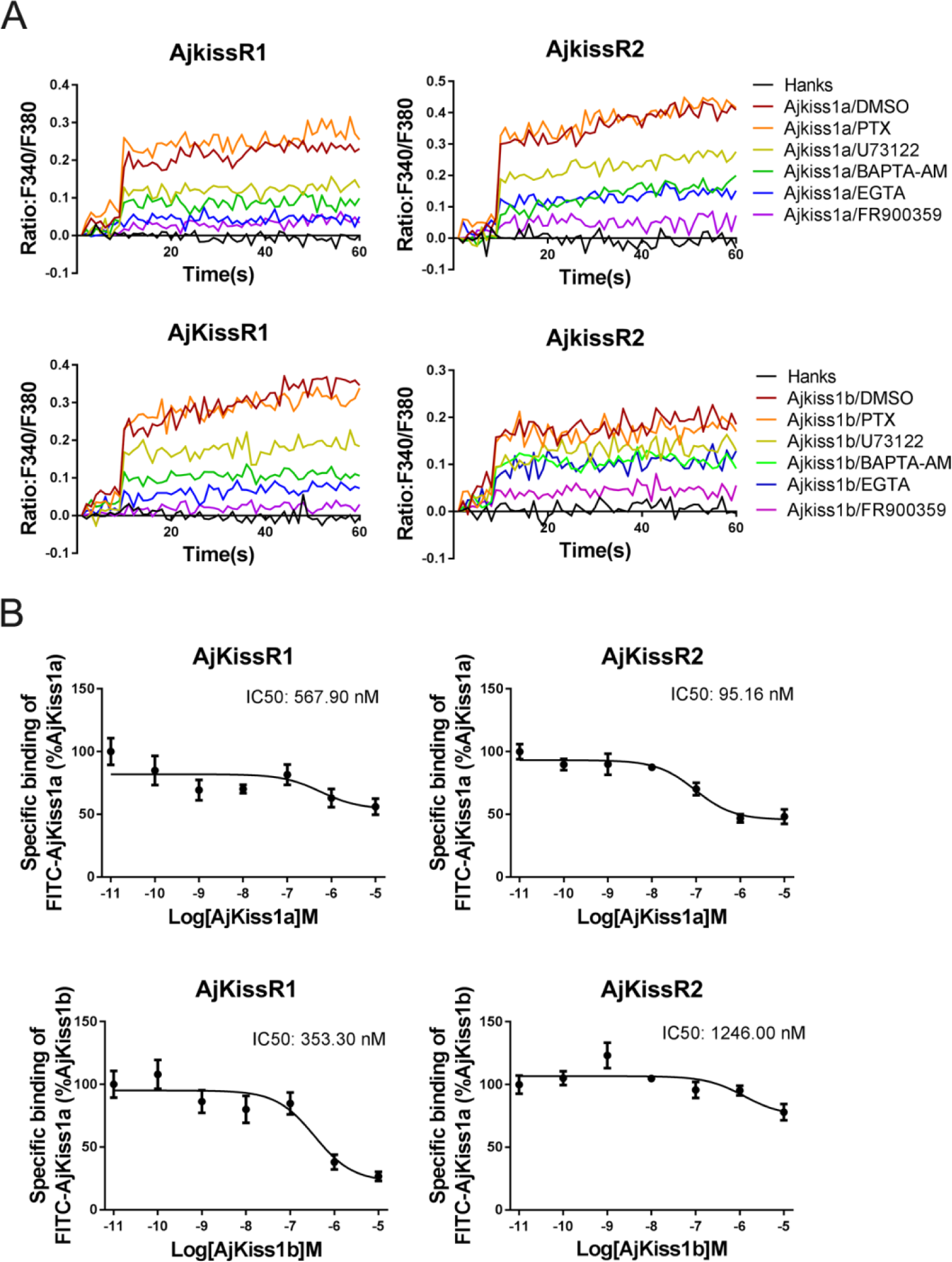
*Apostichopus japonicus* kisspeptin (Kp) receptors are directly activated by Kps via a G_αq_-dependent pathway**. A.** Intracellular Ca^2+^ mobilization in AjKissR1 and AjKissR2 expressing HEK293 cells was measured in response to 100 nM AjKiss1a or AjKiss1b pretreated with DMSO, G_αq_ protein inhibitor (FR900359, 1.0 μM), PLC inhibitor (U73122, 1.0 μM), intracellular calcium chelator (BAPTA-AM, 100.0 μM), or extracellular calcium chelator (EGTA, 5.0 mM). **B.** Competitive binding of 1.0 μM FITC-AjKiss1a to AjKissR1 or AjKissR2 in the presence of the indicated concentration of AjKiss1a or AjKiss1b. Error bars represent the SEM for 3 independent experiments.

### Functional expression of putative Kp receptors

To verify the exact expression and localization of the putative *A. japonicus* Kp receptors, AjKissR1 and 2 with an N-terminal FLAG-tag or with enhanced green fluorescent protein (EGFP) fused to the C-terminal end, were constructed and stably or transiently expressed in human embryonic kidney 293 (HEK293) cells. As shown in Fig. 2A, confocal microscopy revealed that AjKissR1 and 2 were predominantly expressed and localized to the cell surface, with some intracellular accumulation, in the absence of the ligand in HEK293 cells. Next, to examine whether AjKissR1 and AjKissR2 are activated by synthetic Kps, the calcium probe fura-2-based Ca^2+^ mobilization assay was performed. As shown in Fig. 2B, both AjKiss1a and AjKiss1b elicited a rapid increase of intracellular Ca^2+^, in a concentration-dependent manner, in HEK293 cells transfected with AjKissR1 and AjKissR2, respectively. However, AjKissR1 was preferentially activated by AjKiss1b, with an EC50 value of 8.06 nM (Fig. 2B2), whereas AjKissR2 was more specifically activated by AjKiss1a, with an EC50 value of 1.98 nM (Fig. 2B1).

Agonist-mediated internalization from the cell surface to the cytoplasm has been recognized as a key mechanism in regulating the strength and duration of GPCR-mediated cell signaling and to directly reflect the activation of the receptor [32, 33]. In this study, C-terminal fusion expression of AjKissR1 and 2 with EGFP was used to track the internalization and trafficking of receptors. As shown in Fig. 2C, AjKissR1 and 2 were activated by AjKiss1b and AjKiss1a, respectively, to undergo significant internalization from the plasma membrane to the cytoplasm. These data provide clear evidence that AjKissR1 and 2 are functional receptors that are specific for neuropeptides AjKiss1b and AjKiss1a, respectively.

### Ligand selectivity of *A. japonicus* Kp receptors

To examine the cross-reactivity of *A. japonicas* and vertebrate Kp receptors, *A. japonicus*, human, frog, and zebrafish Kps (hKiss1-10, XtKiss1b-10, zfKiss1-10, and zfKiss2-10) were used to detect their potential in triggering intracellular Ca^2+^ mobilization. As indicated in Fig. 2, for AjKissR1, hKiss1-10 and XtKiss1b-10 exhibited higher potency, however, both zfKiss1-10 and zfKiss2-10 showed much lower potency in eliciting Ca^2+^ mobilization (Fig. 3A), while for the activation of AjKissR2, XtKiss1b-10, zfKiss1-10, and zfKiss2-10 had a higher potency than hKiss1-10 (Fig. 3B). However, human neuropeptide S (NPS) showed no potency for the activation of both AjKissR1 and AjKissR2 (Fig. 3C and D). Further analysis demonstrated that both AjKiss1a and AjKiss1b could activate hKiss1R, zfKiss1Ra, and zfKiss1Rb with different potency (Fig. 3E, F and G).

### *A. japonicus* Kp receptors are directly activated by Kps via a G_αq_-dependent pathway

A previous study has demonstrated that in mammals, Kiss1R couples to G_αq_ protein, triggering PLC, intracellular Ca^2+^ mobilization, and the PKC signaling cascade in response to agonists [34]. To elucidate G protein coupling in the activation of both AjKiss1a and AjKiss1b, a combination of functional assays, with different inhibitors, was performed. As shown in Fig. 4A, AjKiss1a and AjKiss1b-eliciting Ca^2+^ mobilization through receptors AjKissR1 and AjKissR2, respectively, were completely blocked by pretreatment with FR900359, a specific inhibitor of G_αq_ protein [35], and also significantly attenuated by PLC inhibitor U73122, extracellular calcium chelator EGTA, and intracellular calcium chelator 1,2-bis(o-aminophenoxy)ethane N,N,N’,N’-tetraacetic acid acetoxymethyl ester (BAPTA-AM) [36].

Next, a competitive binding assay was established by using a synthesized FITC-tagged AjKiss1a at the N-terminus (FITC-AjKiss1a), for assessing the direct interaction of AjKissR1 and AjKissR2 with AjKiss1a and AjKiss1b. Functional assays revealed that FITC-AjKiss1a exhibited the potential to induce Ca^2+^ mobilization comparable to the wild-type neuropeptide (Figure 4–figure supplement 1). The competitive displacement of FITC-AjKiss1a with AjKiss1a and AjKiss1b in HEK293/AjKissR1 and AjKissR2 cells was measured by FACS (Fluorescent Activated Cell Sorting) analysis. As shown in Fig. 4B, unlabeled AjKiss1a and AjKiss1b were found to compete with FITC-labeled AjKiss1a with IC_50_ values of 95.16 and 353.30 nM in AjKissR2 and AjKissR1-transfected HEK293 cells, respectively.

### AjKissR1 and AjKissR2 are activated by AjKiss1b-10 and signal through the G_αq_-dependent MAPK pathway

Since AjKiss1b-10 exhibited high potency to activate both AjKissR1 and AjKissR2 in HEK293 cells (Figure 5–figure supplement 1), it was used to conduct further *in vitro* and *in vivo* experiments. The previous results reveal that the AjKissR1 and AjKissR2 can be activated by ligands and signals through G_αq_-dependent Ca^2+^ mobilization; however, the detailed signaling pathway remained to be elucidated. To address this and to evaluate AjKissR1 and AjKissR2 mediated signaling pathway, different inhibitors were used to test intracellular ERK1/2 activation in 293 cells, expressing AjKissR1 and AjKissR2, treated with Ajkiss1b-10. As shown in Fig. 5A, stimulation with AjKiss1b-10, led to the activation of both AjKissR1 and AjKissR2, inducing significant ERK1/2 activation. Further assessment demonstrated that AjKissR1 or AjKissR2-mediated activation of ERK1/2 was significantly blocked by the PLC inhibitor, u73122 (10 μM), and the PKC inhibitor, Gö6983 (1 μM) (Fig. 5B and C). Moreover, we determined that PKCα, PKCβI, and PKCβII are involved in the activation of the MAPK pathway, using a PKC subtype recruitment assay (Fig. 5D and E). Overall, these results suggest that AjKissR1 and AjKissR2, once activated by ligand, can activate the MAPK cascade, particularly ERK1/2, via the G_αq_/PLC/PKC signaling pathway (Fig. 5F).

**Figure 5.**
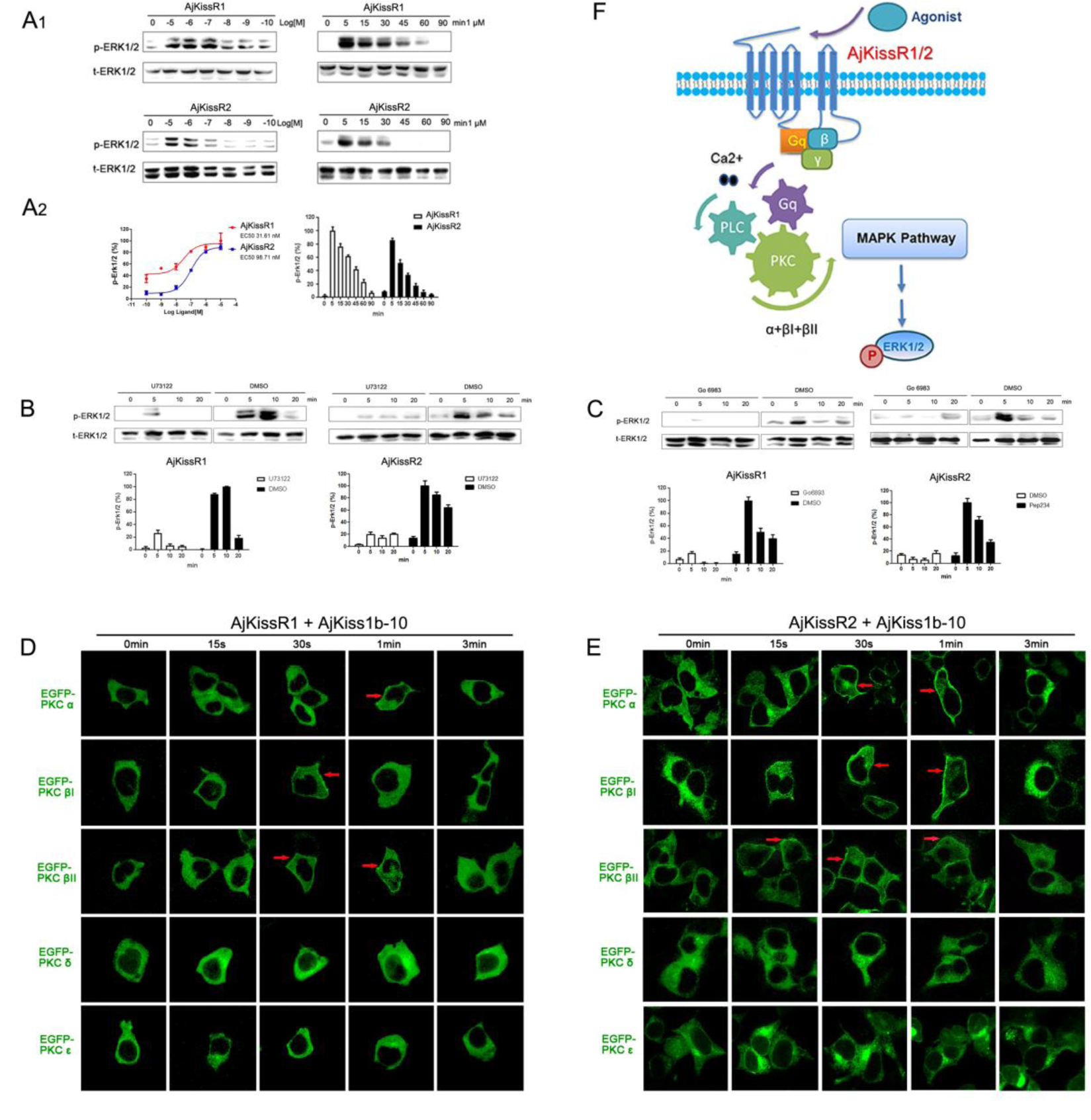
Cell signaling pathway mediated by AjKissR1 or AjKissR2. **A.** Concentration-dependence and time course of AjKiss1b-10 stimulated phosphorylation of ERK1/2 in stable FLAG-AjKissR1 or FLAG-AjKissR2-expressing HEK293 cells, which were incubated with indicated concentrations or times. **B-C.** ERK1/2 phosphorylation, mediated by AjKiss1b-10, was blocked in FLAG-AjKissR1 or FLAG-AjKissR2-expressing HEK293 cells, pretreated with PLC or PKC inhibitor. Serum-starved HEK293 cells were pretreated with DMSO, PLC inhibitor (U73122, 10 μM), or PKC inhibitor (Gö6983, 10 μM). **D-E.** Role of various PKC isoforms in the activated signaling pathways of sea cucumber kisspeptin receptor. HEK293 cells, co-transfected with FLAG-AjKissR1 or FLAG-AjKissR2 and different EGFP-PKC isoforms, were stimulated by 1 μM AjKiss1b-10 for the indicated time and then examined by confocal microscopy. Red arrows denote the recruitment of EGFP-PKC isoforms on cell membrane. **F.** Schematic diagram of agonist-induced *A. japonicus* kisspeptin receptor activation. AjKiss1b-10 binding to AjKissR1 or AjKissR2 activates G_αq_ family of heterotrimeric G protein, which leads to dissociation of the G protein subunits Gβγ, and activates PLC, leading to intracellular Ca^2+^ mobilization, which activates PKC (isoform α and β) and stimulates phosphorylation of ERK1/2. The p-ERK1/2 was normalized to a t-ERK1/2. Error bars represent SEM for 3 independent experiments.

### Physiological functions of the Kp signaling system in *A. japonicus*

To further assess the physiological roles of the Kp signaling system in *A. japonicus*, we examined the tissue distribution of *A. japonicus* Kp/KpR, using custom rabbit polyclonal antibodies for *A. japonicus* kisspeptin precursor and AjKissR1. Tissue-specific western blot analysis revealed the expression of the kisspeptin precursor in the respiratory tree (RET), ovary (OVA), testis (TES), and anterior part (ANP, containing nerve ring as shown in Figure 6–figure supplement 1E, F) of mature sea cucumbers (maturity of gonads was evaluated by H&E staining, as shown in Figure 6–figure supplement 1B). AjKissR1 was detected in the RET, OVA, ANP, and muscle (MUS) (Fig. 6A). To reveal the *in situ* distribution of the kisspeptin precursor and receptor, we performed immunofluorescence labeling on tissue sections. Consistent with results from the western blot assay, significant expression of the kisspeptin precursor was observed in the RET, TES, and nerve ring in ANP sections, with no expression in the OVA and MUS; AjKissR1 expression was observed in the RET, OVA, MUS and nerve ring in ANP sections, with rare expression in TES (Fig. 6B). At the cellular level, the kisspeptin precursor was mainly detected in the coelomic epithelium of RET, while the AjKissR1 was detected in the brown bodies, which can be found in luminal spaces of RET and might be related with foreign material removal [37]. In particular, significant expression and cell membrane localization of AjKissR1 was detected in oocytes, indicating the consistent molecular property of AjKissR1 *in vivo* and *in vitro*. From the TES sections, significant fluorescence signal of the kisspeptin precursor, while a weak signal of AjKissR1, can be detected in spermatogenic epithelium. Significant expression of AjKissR1 was detected in the epithelium of muscle from MUS sections. Moreover, from the ANP sections, the kisspeptin precursor was detectable in the outter surface part of nerve ring (mainly containing the cell body of neurons, as shown in Figure 6–figure supplement 1F2), while the AjKissR1 was detected in the internal region of nerve ring (mainly containing axon of neurons, as shown in Figure 6–figure supplement 1F2).

**Figure 6.**
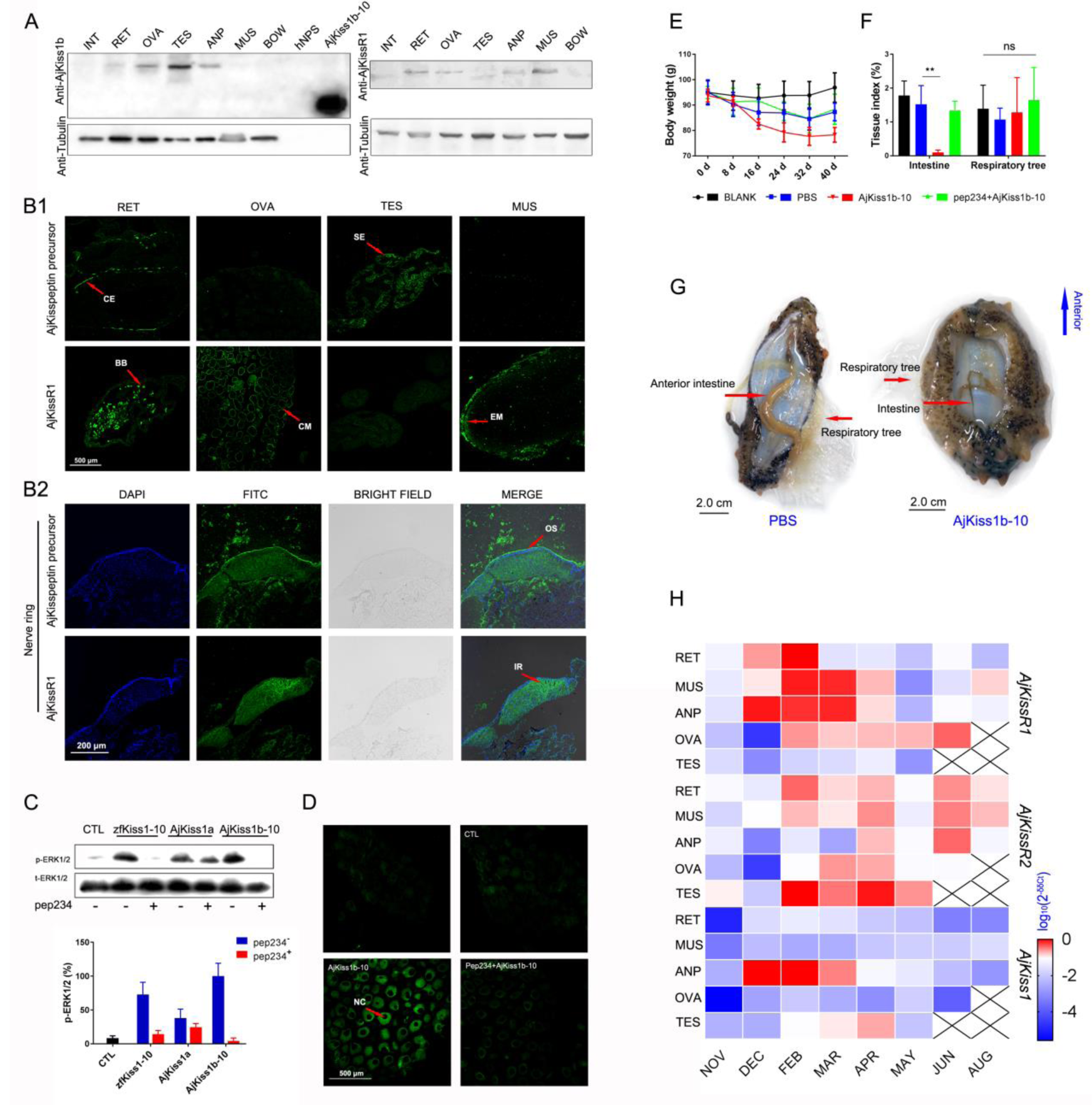
Physiological function analysis of Kp/KpR signaling systems in *Apostichopus japonicus*. **A.** Western Blot analysis of *A. japonicus* kisspeptin precursor and AjKissR1 in different tissues of sea cucumber. (INT) intestine, (RET) respiratory tree, (ANP) anterior part, (OVA) ovary, (TES) testis, (MUS) muscle, and (BOW) body wall. **B.** Immunofluorescence histochemical staining of *A. japonicus* kisspeptin precursor and AjKissR1 in RET, OVA, TES, MUS (B1) and nerve ring (B2) of the sea cucumber. (CE) coelomic epithelium, (BB) brown body, (CM) cell membrane, (SE) spermatogenic epithelium, (EM) epithelium of muscle, (OS) outter surface, (IR) internal region. **C.** ERK1/2 phosphorylation activity of Kps and inhibitory effect of pep234 on the cultured ovary of sea cucumber. Samples were evaluated after 2 h of ligand administration, with or without a 4 h pretreatment of pep234, in optimized L15 medium at 18 °C. Error bars represent SEM for 3 independent experiments. **D.** Immunofluorescence histochemical staining of pERK signal in cultured oocytes of sea cucumber. Samples were collected and fixed after 2 h of ligand administration with or without a 4 h pretreatment of pep234, in optimized L15 medium at 18 °C. NC indicates nucleus of oocytes. **E–F.** Variation of body weight (**E**) and tissue index (**F**) over 40 days of stimuli treatment. Each symbol and vertical bar represent SEM (n=5). * indicates significant differences (P < 0.05) and ** indicates extremely significant differences (P < 0.01), ANOVA, Tukey‟s multiple comparison test. **G.** Degenerated intestine in AjKiss1b-10 treated sea cucumbers. **H.** Heatmap showing the expression profile of *A. japonicus* kisspeptin and kisspeptin receptors (*AjKissR1/R2* and *AjKiss1*) in different tissues and developmental stages of sea cucumber. The variation in color represents the relative expression level of each gene in different samples (normalized against the peak values in all samples and logarithmized). The number of tissues used for all samples is six, except for the number of ovary samples, with one in NOV (November) and JUN (June), three in DEC (December) and FEB (February), five in MAR (March), and six in APR (April) and MAY (May), and in testis, with two in NOV (November) and DEC (December), four in FEB (February) and MAR (March), and six in APR (April) and MAY (May).

To verify the physiological function of *A. japonicus* kisspeptins, cultured oocytes were stimulated by different Kps. As shown in Fig. 6C, significant ERK phosphorylation signal can be detected by western blot assay in different Kp-treated oocytes that can be blocked by kisspeptin antagonist pep234 (1 μM) in zfKiss1-10 or AjKiss1b-10 administrated cells (inhibitory effect of pep234 was preapproved *in vitro* as shown in Figure 6–figure supplement 2). Further detection of the pERK signal in AjKiss1b-10 treated oocytes by confocal microscopy demonstrated the physiological activation of this pathway by AjKiss1b-10 and pep234 on *A. japonicus* cells (Fig. 6D).

Based on the confirmation of their functional activity in cultured oocytes, AjKiss1b-10 and pep234 were used to conduct further *in vivo* experiments. Sea cucumbers treated with AjKiss1b-10 for 40 days exhibited weight loss (p=0.0583, Tukey‟s multiple comparison test, as shown in Fig. 6E) and extremely significant intestinal degeneration (p=0.0001, Tukey‟s multiple comparison test, as shown in Fig. 6F, G), which are the characteristic phenotypes of aestivating *A. japonicus* [38]. Moreover, extremely significant elevation of pyruvate kinase PK transcription (p=0.0001, Tukey‟s multiple comparison test, as shown in Figure 6–figure supplement 3A), which is the rate-limiting enzyme in the regulation of glycolysis and metabolic depression in aestivating *A. japonicus* [39], was detected in the respiratory tree, while a significant decrease of PK transcription was found in muscle (p=0.0497, Tukey‟s multiple comparison test, as shown in Figure 6–figure supplement 3A). To evaluate the potential role of AjKiss1b-10 in regulating reproductive activity, we examined the estradiol (E2) levels in the coelomic fluid of sea cucumber; however, no significant difference was observed in animals treated with AjKiss1b-10 (Figure 6–figure supplement 3B).

The transcriptional expressions of the *A. japonicus* Kp precursor (*AjKiss1*) and Kp receptors (*AjKissR1/2*) were investigated at different stages of reproductive development using the qPCR method. Two-year old sea cucumbers, with 85.29±9.47 g body weight (Figure 6–figure supplement 4A), were collected and various tissues were sampled for further analysis. As shown in Figure 6–figure supplement 4B, notable changes in the relative gut mass and the relative ovary weight of sea cucumber were detected in the developing reproductive stage from November to April, mature reproductive stage in May, after spawning in June, and during aestivation in August. At all stages, *AjKissR1/2* expression was detectable in the majority of sea cucumber tissues, especially after February (Fig. 6H), while significant expression of *AjKiss1* was found in the ANP from December to April with a peak value detected in February. Taken together, the high expression levels of *A. japonicus* Kp precursor mRNA during reproductive development suggests its role in the regulation of reproduction, while the wide distribution of *AjKissR1* and *AjKissR2,* in the other tissues investigated, indicates diverse functions for these two receptors.

## Discussion

The functional characterization of neuropeptides or secretory neurons of non-vertebrates contributes to our understanding of the evolutionary origin and conserved roles of the neurosecretory system in animals, especially in Ambulacrarians (deuterostomian invertebrates including hemichordates and echinoderms), which are closely related to chordates [3, 8]. The hypothalamic neuropeptide kisspeptin (Kp), acts as a neurohormone and plays important roles in the regulation of diverse physiological processes in vertebrates, including reproductive development [40, 41], metastasis suppression [42], metabolism and development [43-45], behavioral and emotional control [46], and the innate immune response [47].

Though a functional Kp/KpR system has been demonstrated in the chordate amphioxus and a number of invertebrate Kp/KpR genes have been predicted recently, missing experimental identification of a Kp-type system in non-chordates makes it difficult to determine if this signaling system has an ancient evolutionary origin in invertebrates or if it evolved *de novo* in the chordate/vertebrate lineages. In this study, two Kp receptors from the sea cucumber *A. japonicus*, AjKissR1 and AjKissR2, have been established to have a high affinity for synthetic Kps from *A. japonicus* or vertebrates and to share similar intracellular signaling, via the G_αq_/PLC/PKC/MAPK pathway. Results from the *in vivo* investigation indicate that the Kp/KpR system in sea cucumber might be involved in both metabolic and reproductive control. Given the highly conserved intracellular signaling pathway and physiological functions revealed for the *A. japonicus* Kp/KpR system, it is interesting to speculate that Kp signaling might have originated from non-chordate invertebrates.

### Two putative Kp receptors can be activated by multiple synthetic Kp-type peptides in *A. japonicus*

Kps or KpRs in Chordata have been functionally recognized in various species. Virtual screening of the transcriptome and genome sequence data for neuropeptide precursors has made a great contribution to Kp/KpR paralogous gene prediction in Ambulacrarians and provides valuable infromation for further investigation (Fig. 7). In 2013, Kp-type receptors were first annotated in the genome of the acorn worm (*S. kowalevskii*) and purple sea urchin (*S. purpuratus*) [20, 48]. Moreover, a Kp-type neuropeptide precursor with 149-amino acid residues was identified in the starfish *A. rubens*, comprising two putative Kp-type peptides, ArKp1 and ArKp2 [24]. Subsequently, *in silico* analysis of neural and gonadal transcriptomes enabled the virtual discovery of Kps in the sea cucumbers *H. scabra* and *H. glaberrima* [23]. Moreover, the presence of Kps in extracts of radial nerve cords was confirmed by proteomic mass spectrometry in the crown-of-thorns starfish *A. planci* [49]. Recently, a 180-residue protein comprising two putative Kp-type peptides has been predicted and a C-terminally amidated peptide GRQPNRNAHYRTLPF-NH2 was confirmed by mass spectrometric analysis of centrol nerve ring extracts [25]. These advances provide a basis for experimental studies on the Kp/KpR system in echinoderms.

**Figure 7.**
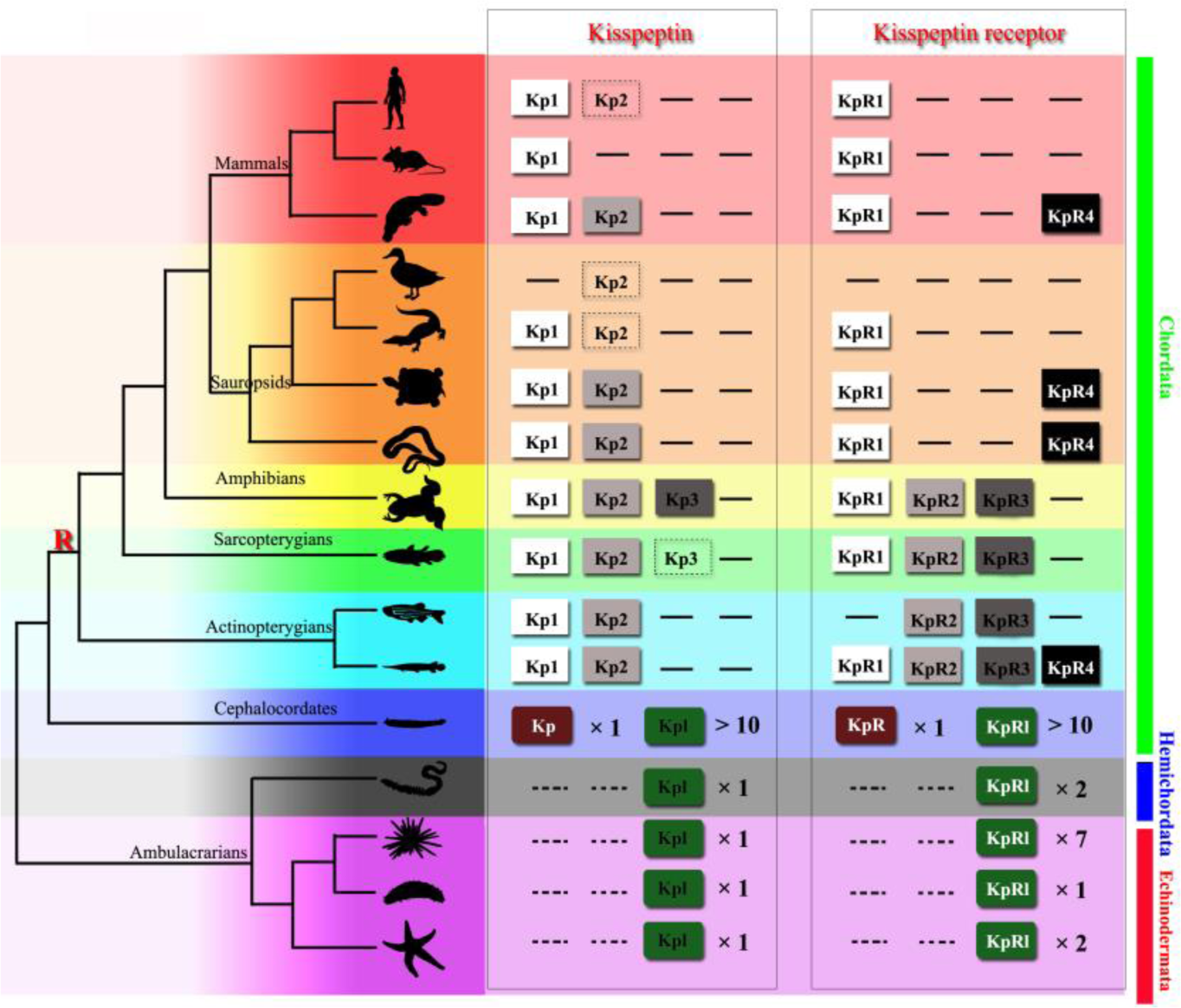
Recently identified Kisspeptin (Kp) or Kisspeptin receptor (KpR) genes among some deuterostomes. The species, indicated by silhouette images downloaded from the *PhyloPic* database, were clustered in a phylogenetic tree and classified by different colors. Red highlighted “R” indicates a whole-genome duplication event. Kp/KpR indicates the identified Kisspeptin/Kisspeptin receptor gene, and Kpl/KpRl indicates predicted Kisspeptin-like/Kisspeptin-like receptor gene. Dashed boxes denote symbols indicate pseudogenes. Arabic numerals indicate the number of genes identified or predicted from public data. The evolutionary tree of indicated species was modified from Pasquier et al. 2014 [18]. Image credits: All silhouettes from PhyloPic, human by T. Michael Keeseyacorn; mouse by Anthony Caravaggi; platypus by Sarah Werning; duck bySharon Wegner-Larsen; crocodile by B Kimmel; turtle by Roberto Díaz Sibaja; python by V. Deepak; frog uncredited; coelacanth by Yan Wong; zebra fish by Jake Warner; spotted gar by Milton Tan; Branchiostoma by Mali’o Kodis, photograph by Hans Hillewaert; acorn worm by Mali’o Kodis, drawing by Manvir Singh; starfish by Hans Hillewaert and T. Michael Keesey; sea cucumber by Lauren Sumner-Rooney; sea urchin by Jake Warner;

In the present study, we cloned the full length of *Kiss* cDNA sequence from the nerve ring, encoding a putative Kp precursor, which has been predicted from the proteomic analysis of *A. japonicas* [25] and synthesized the peptides AjKiss1a (32aa), AjKiss1a-15, AjKiss1a-13, AjKiss1a-10, AjKiss1a (18aa), and AjKiss1b-10, for further experimental tests. Two candidate *A. japonicus* Kp receptors were screened from genomic data, based on the sequence of the identified kisspeptin receptors [19, 20, 29-31, 48, 50] and functionally characterized. Our data shows that despite a low percentage homology between AjKissR1 and 2, synthetic *A. japonicus* Kp peptides (AjKiss1a and AjKiss1b) could activate both the receptors, thereby initiating significant receptor internalization and extensive Ca^2+^ mobilization, albeit with a different potency. This is consistent with previous studies demonstrating that in non-mammalian species, synthetic Kiss1 and Kiss2 activated Kp receptors *in vitro* with differential ligand selectivity [51, 52]. In particular, the truncated peptide AjKiss1b-10 demonstrated high activity to elicit intracellular Ca^2+^ mobilization in AjKissR1/2 expressing HEK293 cells, while the truncated peptides, AjKiss1a-15, AjKiss1a-13, and AjKiss1a-10, failed to activate the receptors. The functional activity of the truncated peptide AjKiss1b-10 is not unusual, considering that alternative cleavage occurs in the Kps of verterbrates [51, 53]; however, the inactivity of AjKiss1a-15, which was identified from mass spectrometric detection in *A. japonicus* [25], raised more questions about the functional and structural characteristics of this neuropeptide and requires further investigation.

### Cross interaction between *A. japonicus* and the Kp/KpR systems of vertebrates confirmed the existence of Kp signaling systems in Echinoderm

In the mammalian genome, a single *Kiss1* gene produces a mature 54-amino acid peptide, Kp-54, which is further proteolytically truncated to 14 and 13 amino acid carboxyl-terminal peptides, Kp-14 and Kp-13, with a common C-terminal decapeptide (Kp-10) core [53, 54]. In non-mammalian vertebrates, two paralogous Kp genes, *Kiss1* and *Kiss2*, are present in the genome of teleosts, producing two mature peptides, which share the highly conserved Kp-10 region with mammalian Kps [50, 51, 55]. Unlike mammalian and non-mammalian vertebrates, in the sea cucumber *A. japonicas*, only one Kp gene was annotated and isolated. However, sequence analysis revealed that the Kp gene encodes a 180 amino acid peptide precursor, which is proteolytically cleaved to two mature peptides, consistent with other KPs identified in the phylum Echinodermata [22, 24, 49]. Both putative mature peptides have a C-terminal Leu-Pro-Phe-amide motif, instead of the Arg-Phe-amide motif common in vertebrate Kps, and exhibit a much lower identity with vertebrate Kp sequences. Thus, the experimental evidence collected from functional interaction studies, between *A. japonicus* Kps and KpRs, was not sufficient to support a definite relationship between the neuropeptide and the receptor.

To address this issue, the cross interaction between vertebrate and *A. japonicus* Kp/KpR was evaluated in this study. Our specificity analysis showed that human, frog, and zebrafish KPs, hKiss1-10, XtKiss1b-10, and zfKiss1-10 and zfKiss2-10, were potent in activating both AjKissR1 and AjKissR2, while the human neuropeptide S (hNPS, as a negative control) showed no potency for the activation to AjKissR1 nor AjKissR2. Likewise, neuropeptides AjKiss1a and AjKiss1b could potentiate Ca^2+^ signaling by binding the human Kp receptor hKiss1R and zebrafish Kp receptors zfKiss1Ra/b, similar to the corresponding active decapeptides. This, to our knowledge, is the first experimental data directly confirming the connection between the Kp/KpR systems of vertebrates and *A. japonicus*, therefore proving the existence of this neuropeptide system in non-chordate species. Considering the high conservation of the neuropeptides in different echinoderms [4, 22], our finding that the Kp signaling system exists in *A. japonicus* may be extend to other taxa in this phylum.

### Conserved G_αq_/PLC/PKC/MAPK intracellular pathway mediated by *A. japonicus* Kp/KpR system provides insights into the evolution of Kp signaling

It is well established that in mammals, Kiss1R is a typical G_αq_-coupled receptor, triggering PLC, intracellular Ca^2+^ mobilization, and the PKC signaling cascade in response to agonists [16]. However, accumulating evidence shows that in teleosts, while both Kp receptors preferentially activate the G_αq_-dependent PKC pathway, one of them is also capable of triggering the G_αs_-dependent PKA cascade in response to Kp challenge [50, 52]. Using CRE-Luc and SRE-Luc reporting assays, which helps discriminate between the AC/PKA and PLC/PKC signaling pathways, an amphioxus Kp receptor was shown to trigger significant PKC and not PKA signaling, when stimulated by two Kp-type peptides [19].

In this study, our data showed that upon synthetic peptide stimulation, both AjKissR1 and AjKissR2 induced a rapid and transient rise of intracellular Ca^2+^, in a dose-dependent manner, via the G_αq_-coupled signaling pathway. Further investigation of AjKissR1 and AjKissR2 mediated cell signaling indicated that AjKissR1 and AjKissR2 share similar intracellular signaling pathways, via G_αq_/PLC/PKC and ERK1/2 phosphorylation. Our results showed no significant accumulation of cAMP, as detected by ELISA, indicating that G_αs_-dependent PKA signaling was not activated by the Kp receptors of *A. japonicus*. Since the G_αq_-coupled PKC signaling pathway, mediated by identified Kp systems, is conserved in all chordate species and *A. japonicus*, and the G_αs_-dependent PKA signaling was conserved in only a few teleost Kp receptors (mainly from the KpR3 subfamily), we propose that G_αq_-coupled signaling activation originally evolved in this hypothalamic neuropeptide system.

### Reproductive and metabolic regulatory functions identified in *A. japonicus* revealed the ancient physiological roles of the Kp system

Diverse physiological functions of the Kp system have been reported in vertebrate species. In mammals, it is widely established that the Kp signaling system is essential for HPG axis regulation, leading to reproductive control, and the hypothalamic kisspeptin neurons have been found to stimulate pituitary gonadotropin-releasing hormone neurons, which express the kisspeptin receptor, providing a neural pathway of mammalian Kp neuronal system [56]. In non-mammalian species, especially in teleosts, the reproductive function of the Kp system is still controversial, considering the normal reproductive phenotypes observed in fishes in the absence of Kps. A new theory has been proposed that the nonreproductive functions outside HPG regulation, are the conserved roles of Kps in vertebrates [57, 58]. Here, we applied multiple approaches to analyze the potential functions of the recently identified Kp in *A. japonicus*, aiming to give some insights into the ancient physiological roles of the Kp system. As described in this study, the expressional distribution of the *A. japonicus* Kp/KpR protein in multiple tissues suggests the involvement of the Kp signaling system in both reproductive and non-reproductive functions. Interestingly, the unequally expressed Kp/KpR protein levels in gonads, comparatively high Kp precursor level in testis, and high KpR protein levels in ovary demonstrated in our study, revealed differential functions of the Kp system in different genders of sea cucumber. Further investigation from both *in vivo* and *in vitro* experiments would indicate a role for the Kp signaling system in regulating gut function in sea cucumber. Combining the feeding regulatory function of VP/OT-type neuropeptides characterized in echinoderm [8] and the interaction between Kp and VP/OT neural systems [58-60], we suggest that Kp regulation on VP/OT system may exist in echinoderms, requiring further exploration on the possible interaction between these two systems and an evolutionarily conserved function of the Kp system.

## Materials and Methods

### Materials

For cDNA cloning and gene expression analysis in various tissues, individuals of the sea cucumber *A. japonicus* were collected from separate culture ponds in Qingdao (Shandong, China, in 2016–2017). Each batch was acclimated in seawater aquaria (salinity range: 32.21–34.13) for seven days and further dissected, sampled, and stored in liquid nitrogen for future use or directly used for tissue culture. Individuals for *in vivo* experiments (94 ± 4.3 g) were collected from the same culture pond in November 2017, kept in a 500 L tank, and fed with a formulated diet (45% marine mud, 50% Sargasso, and 5% shrimp shell powder) before chemicals were administered. After 15 days, sea cucumbers were randomly assigned to different groups (10 individuals per group). AjKiss1b-10 was dissolved in PBS and intraperitoneal injection of 100 μL AjKiss1b-10 (concentration of 0.5 mg/mL diluted in PBS) or PBS alone, was conducted once every two days, at noon. After 40 days (December 10, 2017 to January 18, 2018) of chemical administration, animals were dissected and the respiratory tree, intestine, muscle, and anterior part tissues were taken as sample from five individuals, for each group and stored in liquid nitrogen for future use. Coelom fluid was collected and stored at **−**20 °C for E2 detection. This experiment was carried out on Xixuan Fishery Technology Island without temperature or light control (sea water temperature 11.5–7.0 °C). Individuals used in the *in vitro* experiments (89 ± 2.4 g) were collected from the same culture pond in May 2017 and the respiratory tree, muscle, body wall, intestine, anterior part (containing nerve ring), and ovary were dissected and further restored in **−**20 °C for western blotting or washed with PBS three times, in aseptic conditions, for tissue culture and *in vitro* experiments.

### Bioinformatic searches and tools

The cDNA sequences were used to query known sequences in GenBank using the blastx utility, BLASTX 2.8.0+ (http://blast.ncbi.nlm.nih.gov/). The cDNA sequence of *A. japonicus* Kp receptors or Kp precursor was translated into the predicted amino acid sequence with DNAMAN 8.0. Analysis of physicochemical properties of proteins was based on Protparam (http://www.expasy.org/tools/protparam.html). Analysis of transmembrane regions in the protein was achieved by TMHMM (http://topcons.cbr.su.se/). The deduced amino acid sequences were aligned using ClustalW. Color align property was generated by the Sequence Manipulation Suite (http://www.bioinformatics.org/sms2/color_align_prop.html). Signal peptide was predicted by SignalP-5.0 Server (http://www.cbs.dtu.dk/services/SignalP/). Phylogenetic tree construction was based on the Maximum Likelihood (ML) Method of Molecular Evolutionary Genetics Analysis (MEGA 5.1). The bootstrap value was repeated 1, 000 times to obtain the confidence value for the analysis.

### Molecular Cloning and Plasmid Construction

To construct the AjKissR1/2 fusion expression plasmids, RT-PCR was performed using total RNA extracted from *A. japonicus* ovaries, to synthesize template cDNA. PCR amplification for coding sequences of *AjKissR1/2* was performed using specific primers, with restriction sites (Supplementary Table 2). The corresponding PCR products were then cloned to pCMV-FLAG and pEGFP-N1 vectors, respectively, using restriction enzymes and Rapid DNA Ligation Kit (Beyotime, China). FLAG-hKiss1R plasmid was constructed using total synthesized DNA (Wuhan Transduction Bio) with specific primers containing restriction sites (Supplementary Table 2). All constructs were sequenced to verify the correct sequences, orientations, and reading frames.

### Cell culture and transfection

HEK293 cells were cultured in DMEM (HyClone) supplemented with 10% FBS, 100 U/mL penicillin, 100 mg/mL streptomycin and 4.0 mM L-glutamine (Thermo Fisher Scientific) at 37 °C in a humidified incubator containing 5% CO_2_. The plasmid constructs were transfected into HEK293 cells by using X-tremeGENE HP (Roche), according to the manufacturer’s instructions. Two days after transfection, stably expressing cells were selected by the addition of 800 mg/L G418.

### Intracellular calcium measurement

The fluorescent Ca^2+^ indicator Fura-2/AM was used to detect intracellular calcium flux [61]. Briefly, the AjKissR1 or AjKissR2 expressing HEK293 cells were washed twice with PBS and suspended at 5×10^6^ cells/mL in Hanks‟ balanced salt solution. The cells were then loaded with 3.0 μM Fura-2/AM for 30 min and washed twice in Hanks‟ solution. The cells were then stimulated with the indicated concentrations of different predicted *A. japonicas* Kps or vertebrate Kps. Finally, intracellular calcium flux was measured for 60 s, by the ratio of excitation wavelengths at 340 and 380 nm, using a fluorescence spectrometer (Infinite 200 PRO, Tecan, Männedorf, Switzerland). All the experiments for measuring Ca^2+^ mobilization were repeated independently at least thrice.

### Receptor localization and translocation assay, by confocal microscopy

For the expression and translocation analysis of receptors, HEK293 cells expressing AjKissR1/2-EGFP were seeded onto glass coverslips in 12-well plates, coated with 0.1 mg/mL poly-L-lysine and allowed to attach overnight under normal growth conditions [61]. The cells were washed three times with PBS and further incubated with or without DAPI for several minutes. The translocation of the receptor was measured with 1.0 μM of various stimuli for 30 min. Cells were washed three times with PBS and then fixed with 4% paraformaldehyde in PBS for 10 min at room temperature. Finally, the cells were mounted in mounting reagent (DTT/PBS/glycerol,1:8:2) and visualized by fluorescence microscopy on a Zeiss laser scanning confocal microscope, which was attached to a Zeiss Axiovert 200 microscope and linked to a LSM5 computer system.

### PKC subtype recruitment assay by confocal microscopy

Kisspeptin/GPR54 mediated PKC subtype recruitment assay in AjKissr1/2-expressing HEK293 cells, was done after treatment with 1 μM of different kisspeptins. HEK293 cells co-transfected with FLAG-AjKissR1 or FLAG-AjKissR2 and various PKC-EGFP were stimulated with AjKiss1b-10 (1 μM) for the indicated periods and then examined by confocal microscopy, for fusion protein localization and translocation assay.

#### Antibodies

The primary antibodies used for pERK1/2, ERK1/2, or β-tubulin detection were: rabbit anti-phospho-ERK1/2(Thr^202^/Tyr^204^) antibody (1:2,000; Cell Signaling Technology), rabbit anti-ERK1/2 antibody (1:2,000; Cell Signaling Technology), and beta-tubulin rabbit monoclonal antibody (1:2,000; Beyotime). To examine the *A. japonicus* kisspeptin precursor or AjKissR1 in various tissues of sea cucumber, AjKiss1b-10 or a peptide corresponding to amino acids Ser^150^∼Trp^174^ of AjKissR1, the second intracellular loop, was synthesized and injected into two rabbits, respectively. The polyclonal antibodies, rabbit anti-AjKiss1b-10 (1:1,000) was prepared by ChinaPeptides and anti-AjKissR1 (1:1,000) was prepared by Wuhan Transduction Bio. The secondary antibodies used were, HRP-conjugated goat anti-rabbit IgG (Beyotime) and FITC-conjugated goat anti-rabbit IgG (Beyotime).

#### Protein extraction and western blotting

To examine the phosphorylation of ERK, cells that expressed AjKissr1/2 or other GPR54s, were incubated for the indicated times with different concentrations of kisspeptins [62]. Subsequently, cells were lysed with lysis buffer (Beyotime) that contained protease inhibitor (Roche) at 4 °C for 30 min on a rocker and then scraped. Proteins were then electrophoresed on a 10% SDS polyacrylamide gel and transferred to PVDF membranes. Membranes were blocked with 5% skim milk, then probed with rabbit anti-phospho-RK1/2(Thr^202^/Tyr^204^) antibody (1:2,000; Cell Signaling Technology), followed by detection using HRP-conjugated goat anti-rabbit IgG (Beyotime). Blots were stripped and reprobed by using anti-ERK1/2 antibody (1:2,000; Cell Signaling Technology), as a control for protein loading.

To detect AjKissR1 in different tissues of sea cucumber, the respiratory tree, intestine, muscle, nerve ring, and ovary was sampled and homogenized with lysis buffer (Beyotime) that contained protease inhibitor (Roche) at 4 °C. Comparable concentrations of proteins were then electrophoresed on a 10% SDS polyacrylamide gel and transferred to PVDF membranes. Membranes were blocked with 5% skim milk, then probed with rabbit anti-AjKissR1 serum (1:1,000), followed by detection using HRP-conjugated goat anti-rabbit IgG (Beyotime). Samples were probed in parallel with anti-tublin antibody (Beyotime), as control for protein loading.

Immunoreactive bands were detected with an enhanced chemiluminescent substrate (Beyotime), and the membrane was scanned by using a Tanon 5200 Chemiluminescent Imaging System (Tanon Science & Technology, Shanghai, China).

#### Ligand competition binding assay

A fluorescence-activated cell sorter (FACS) was used to detect the binding ability of Kps with AjKissR1 or AjKissR2. HEK293 cells, expressing Flag-AjKissR1 or Flag-AjKissR2, were washed with PBS that contained 0.2% bovine serum albumin (FACS buffer). We designed and synthesized N-terminal FITC-labeled AjKiss1a peptides (Supplementary table 1). Different Kps were diluted in the FACS buffer to different concentrations, then added to cells that were incubated on ice for 60–90 min. Cells were washed thrice with the FACS buffer and re-suspended in the FACS buffer with 1% paraformaldehyde, for 15 min. The binding activity of indicated Kp peptides with AjKissR1 or AjKissR2 was determined by measuring the fluorescence of FITC and was presented as a percentage of total binding.

#### Immunofluorescence assay on paraffin-embedded tissue sections

Paraffin sections were baked at 60 °C for 2–4 h and placed in xylene for 15 minutes, twice. The slides were washed twice in 100% ethanol for 10 min each, then in 95% ethanol for 10 min, 85% ethanol for 5 min, 70% ethanol for 5 min, 50% ethanol for 5 min followed by washing with dH_2_O for 5 min, and finally washing with PBS for 5 min. Antigen unmasking was performed in sodium citrate buffer, pH 6, for 10 min at 97 °C and then cooling to room temperature. Endogenous peroxidases were blocked by 10-min incubation in 3.0% hydrogen peroxide. Nonspecific antigens were blocked by a 60-min incubation in 0.3% bovine serum albumin (BSA) in TBST. Slides were incubated with primary antibodies overnight after removing the blocking solution, followed by 2 h incubation with Fluorescein Isothiocyanate (FITC)-conjugated secondary antibodies (FITC-labeled goat anti-rabbit IgG (H+L), Beyotime). Slides was washed with dH_2_O, mounted with antifade mounting medium (Beyotime), and imaged by confocal microscopy.

#### Real-time quantitative PCR (qRT-PCR)

For qRT-PCR, *β-actin* (ACTB) and *β-tubulin* (TUBB) were chosen as the internal control (housekeeping) genes and gene-specific primers were designed based on the ORF sequences [39, 63]. Specific qRT-PCR primers for *AjKissR1/2* and *AjKiss1* were designed based on CDS (Supplementary Table 3). The primers were tested to ensure amplification of single discrete bands, with no primer-dimers. qRT-PCR assays were carried out using the SYBR PrimeScript™ RT reagent Kit (TaKaRa, Kusatsu, Japan) following manufacturer‟s instructions and ABI 7500 Software v2.0.6 (Applied Biosystems, UK). The relative level of gene expression was calculated using the 2^-△Ct^ method and data was normalized by geometric averaging of the internal control genes [64, 65].

#### Tissue culture

For *in vitro* experiments, the ovary and respiratory tree tissues were cut into small pieces of approximately 1 mm^3^ and cultured in Leibovitz L-15 medium (HyClone) supplemented with 12.0 g/L NaCl, 0.32 g/L KCl, 0.36 g/L CaCl_2_, 0.6 g/L Na_2_SO_4_, 2.4 g/L MgCl_2_, 0.6 g/L glucose, 1.5 U/mL penicillin, and 1.5 U/mL streptomycin, at 18 °C in a humidified incubator.

#### Radioimmunoassay

Levels of estradiol (E2) in coelomic fluid or culture medium were measured using the Iodine (^125^I) method [66]. In brief, estradiol levels were measured using Iodine (^125^I) radioimmunoassay kits (Beijing North Institute of Biotechnology, Beijing, China), according to the manufacturer‟s protocol. The binding rate is highly specific with an extremely low cross-reactivity to other naturally occurring steroids, which was less than 0.1% to most circulating steroids.

#### Data Statistics

Statistical analysis was done with GraphPad Prism (version 7.0). Statistical significance was determined using the Student‟s t test and analysis of variance (ANOVA). Probability values that were less than or equal to 0.05 were considered significant (*P < 0.05, **P < 0.01), and all error bars represent standard error of the mean (SEM). All experimental data were gathered from at least 3 independent experiments showing similar results.

## Acknowledgements

The authors of this paper would like to thank Prof. Igor Yu. Dolmatov from National Scientific Center of Marine Biology-Russian Academy of Sciences for his assistance on histomorphological analysis and Prof. Dongdong Xu for his technical assistance and equipment usage. This work was supported by the National Science Foundation of China (Nos. 41876154, 41406137 and 41606150).

## Additional information

## Author contributions

T.W. and N.Z. conceived and coordinated the study. T.W., J.Y., N.Z. and S.G. wrote the main manuscript text, T.W. and X.C. prepared figures 1, 6 and 7, supplementary tables and related supplementary figures, Z.C. and Z.S. prepared figures 2–5, and related supplementary figures. T.W., J.Y., and N.Z. designed the experiments, Z.C., Z.S., Z.Y., K.X., X.X., Q.Y., Y.S., X.C., W.W and Y.T. performed the experiments. T.W., Z.C., Z.S. and N.Z. analyzed the results. L.S., L.Z, S.G. and N.Z. provided technical assistance and expert advice on English writing. All authors reviewed the results and approved the final version of the manuscript.

## Data availability

All the data needed to evaluate the conclusions of the paper are present in the paper and the supplementary information files. All relevant data are available within source data files or from the authors upon reasonable request.

## Supplementary information

Existence and functions of hypothalamic kisspeptin neuropeptide signaling system in a non-chordate deuterostome species This supplementary information section contains the following: Supplementary figures:

**Figure 1–figure supplement 1.**
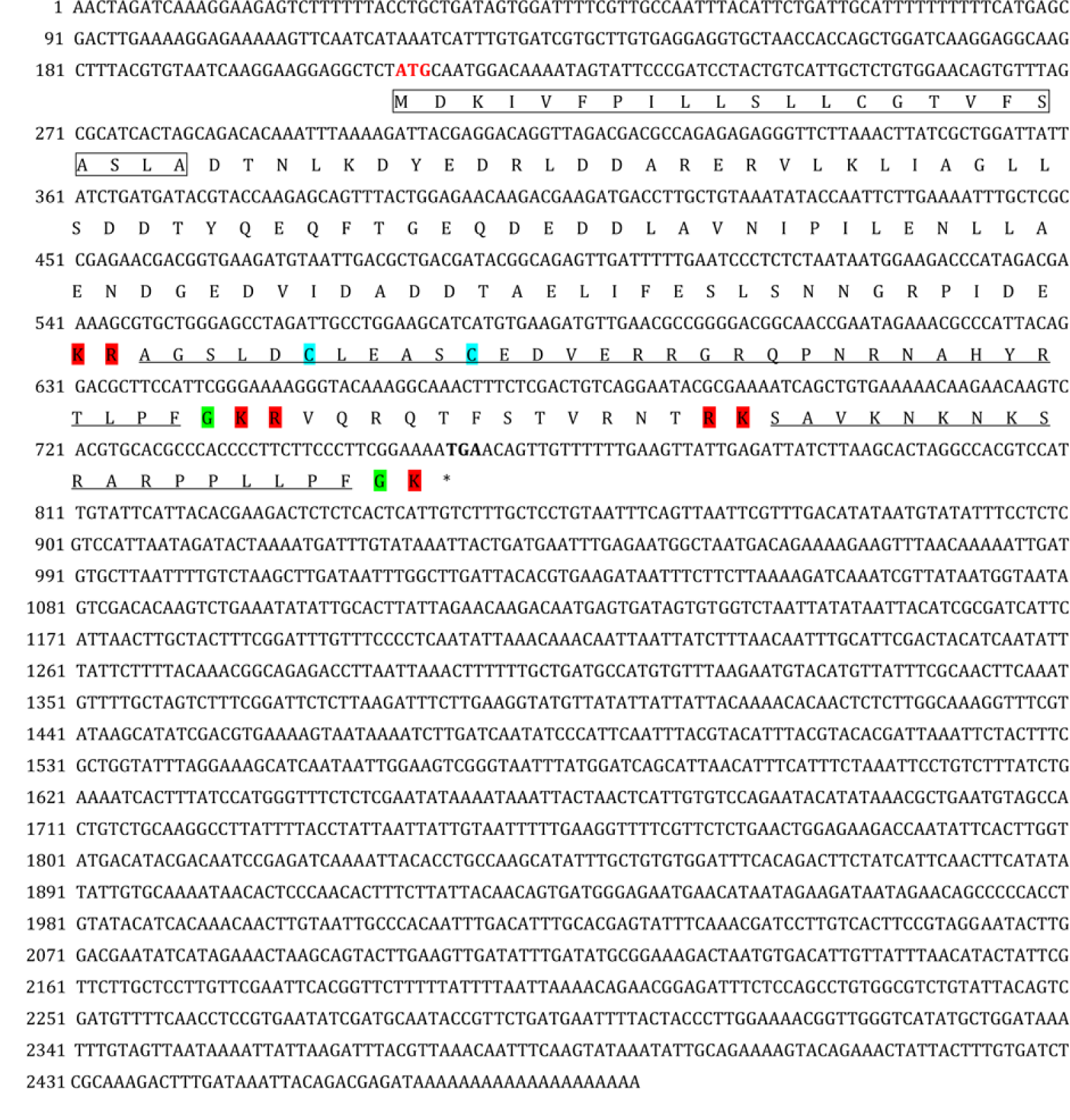
Gene structure of *Apostichopus japonicus* kisspeptin (Kp) precursor. The signal peptide, predicted by online SignalP-5.0 Server, is labeled in box with full lines; the cleavage sites, predicted based on previously known consensus cleavage motifs by using the NeuroPred program, are highlighted in red; glycine residues responsible for C-terminal amidation are highlighted in green; cysteines paired in a disulfide-bonding structure are highlighted in light blue; the predicted mature peptides with C-terminal amidation are noted underlined in black. The initiation codon (ATG) and the termination codon (TGA) are shown in bold.

**Figure 1–figure supplement 2.**
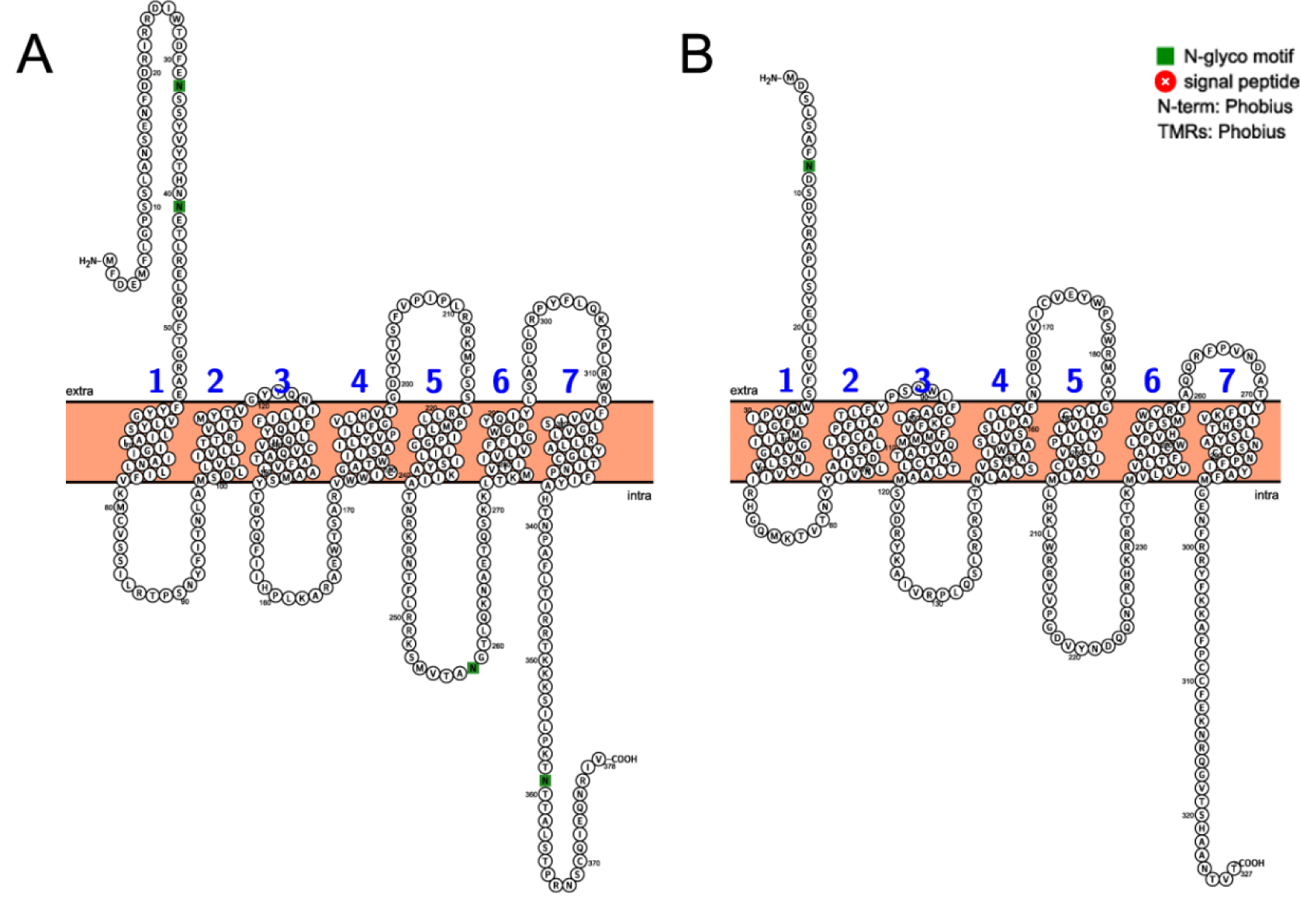
Sequence, topology and annotations of *Apostichopus japonicus* kisspeptin receptors (A: AjKissR1, B: AjKissR2) visualized by a webservice of Protter. http://wlab.ethz.ch/protter/start/.

**Figure 1–figure supplement 3.**
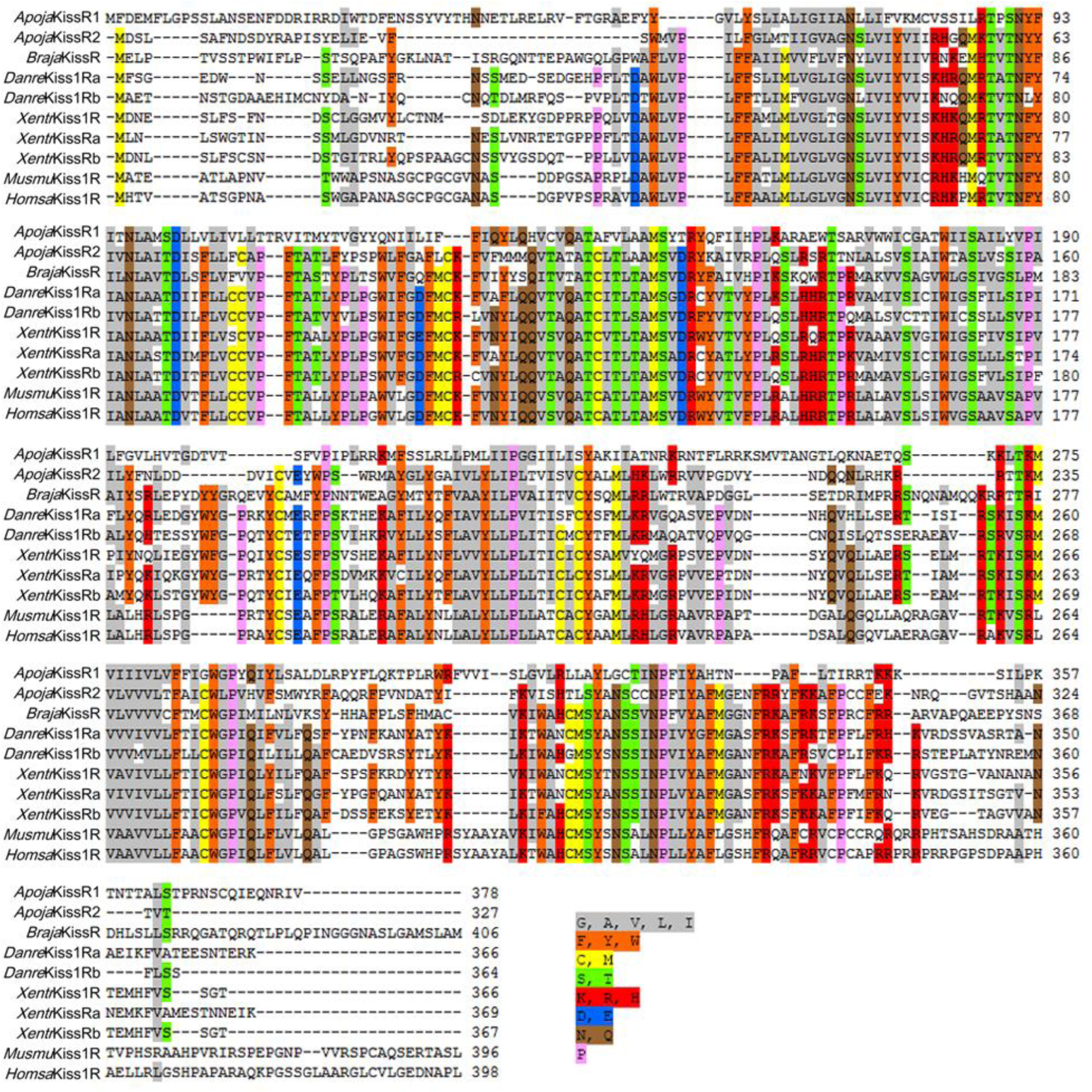
Alignment of the deduced *Apostichopus japonicus* kisspeptin receptor amino acid sequences with functionally characterized chordate GPR54 molecules from other species. Sequences of *Branchiostoma japonicum* kisspeptin (Kp) receptor (*Braja*KissR), *Danio rerio* Kp receptors (*Danre*Kiss1Ra NP_001099149.2 and *Danre*Kiss1Rb NP_001104001.1), *Xenopus tropicalis* Kp receptors (*Xentr*Kiss1R NP_001163985.1, *Xentr*KissRa NP_001165296.1 and *Xentr*KissRb NP_001165295.1), *Mus musculus* Kp receptor (*Musmu*Kiss1R NP_444474.1), and *Homo sapiens* Kp receptor (*Homsa*Kiss1R NP_115940.2) were obtained from GenBank. Alignment was conducted using CLUSTAL W and the color align property was generated using Sequence Manipulation Suite online. Percentage of sequences that must agree for identity or similarity coloring was set as 60%.

**Figure 1–figure supplement 4.**
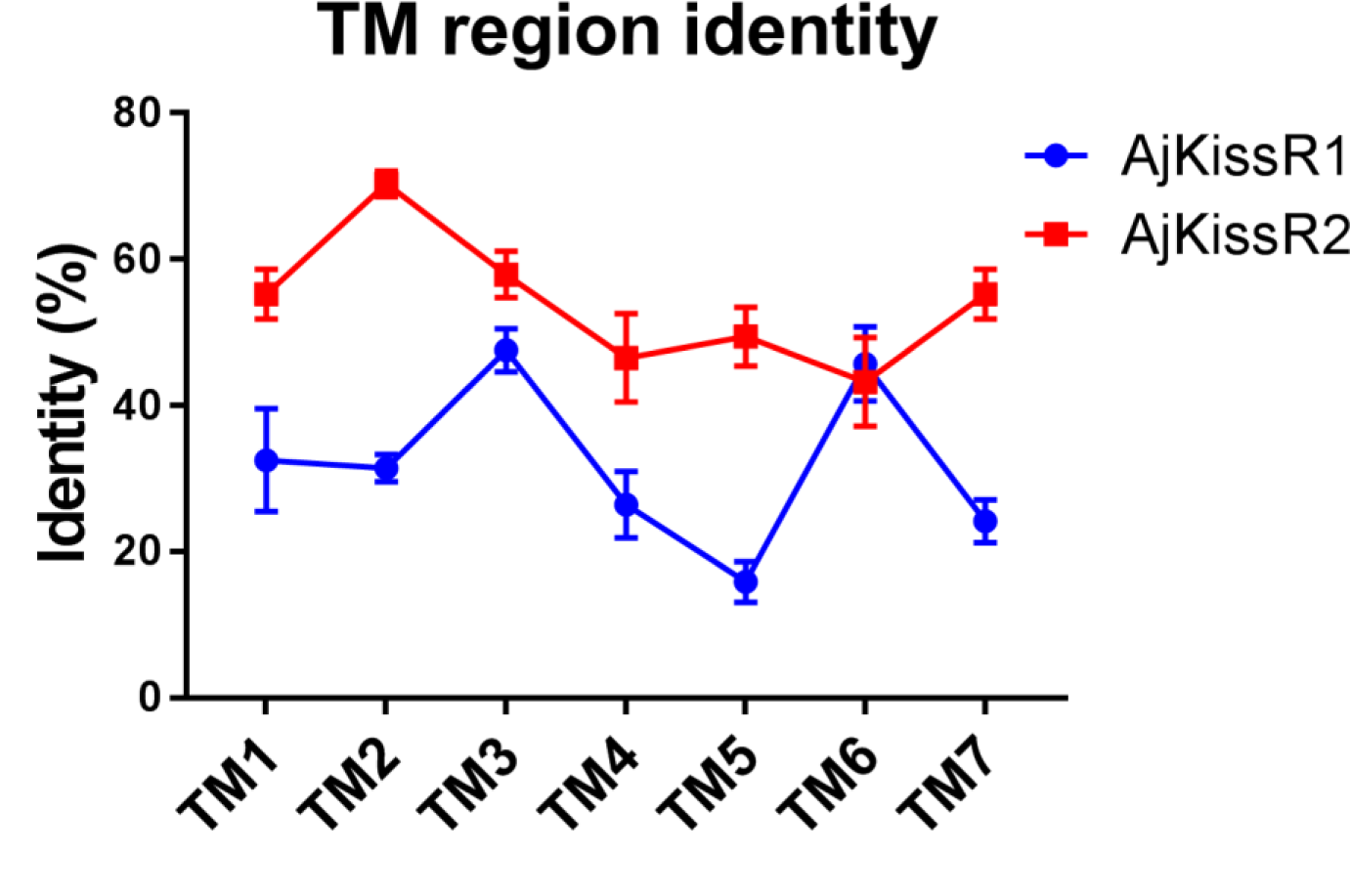
Transmembrane region sequence similarity of *Apostichopus japonicus* kisspeptin receptors to vertebrate kisspeptin receptors. Detailed identities are listed in Figure 1–figure supplement 4 raw data set 1.

**Figure 4–figure supplement 1.**
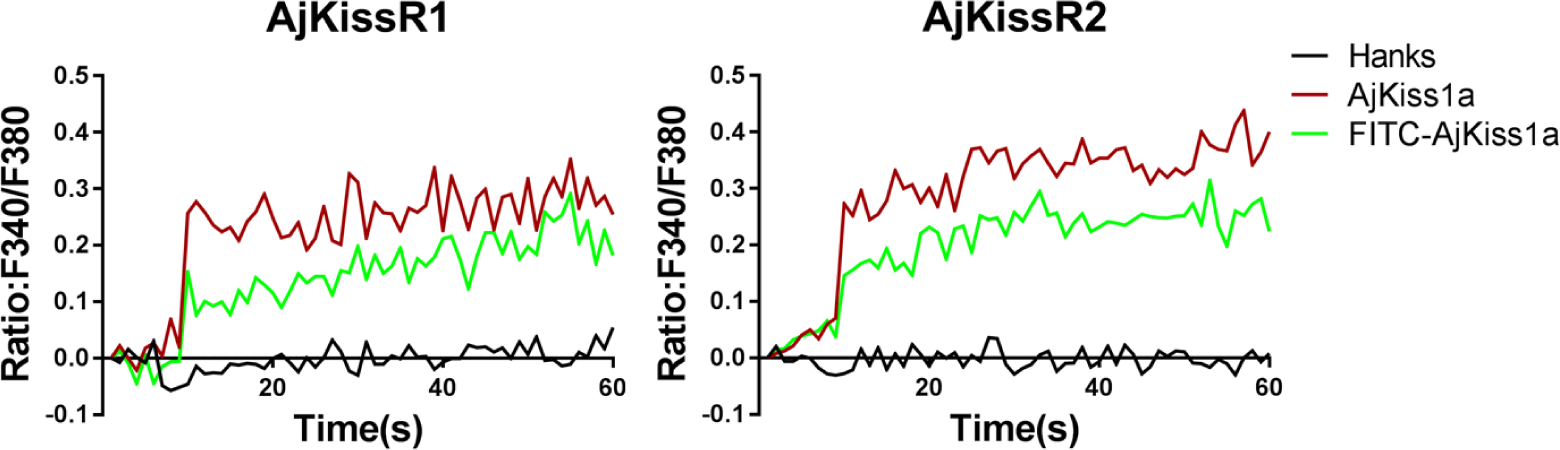
Functional activity of FITC-AjKiss1a evaluated by intracellular Ca^2+^ mobilization detection. Intracellular Ca^2+^ mobilization in AjKissR1/2 expressing HEK293 cells was measured in response to 1.0 μM stimuli using Fura-2/AM.

**Figure 5–figure supplement 1.**
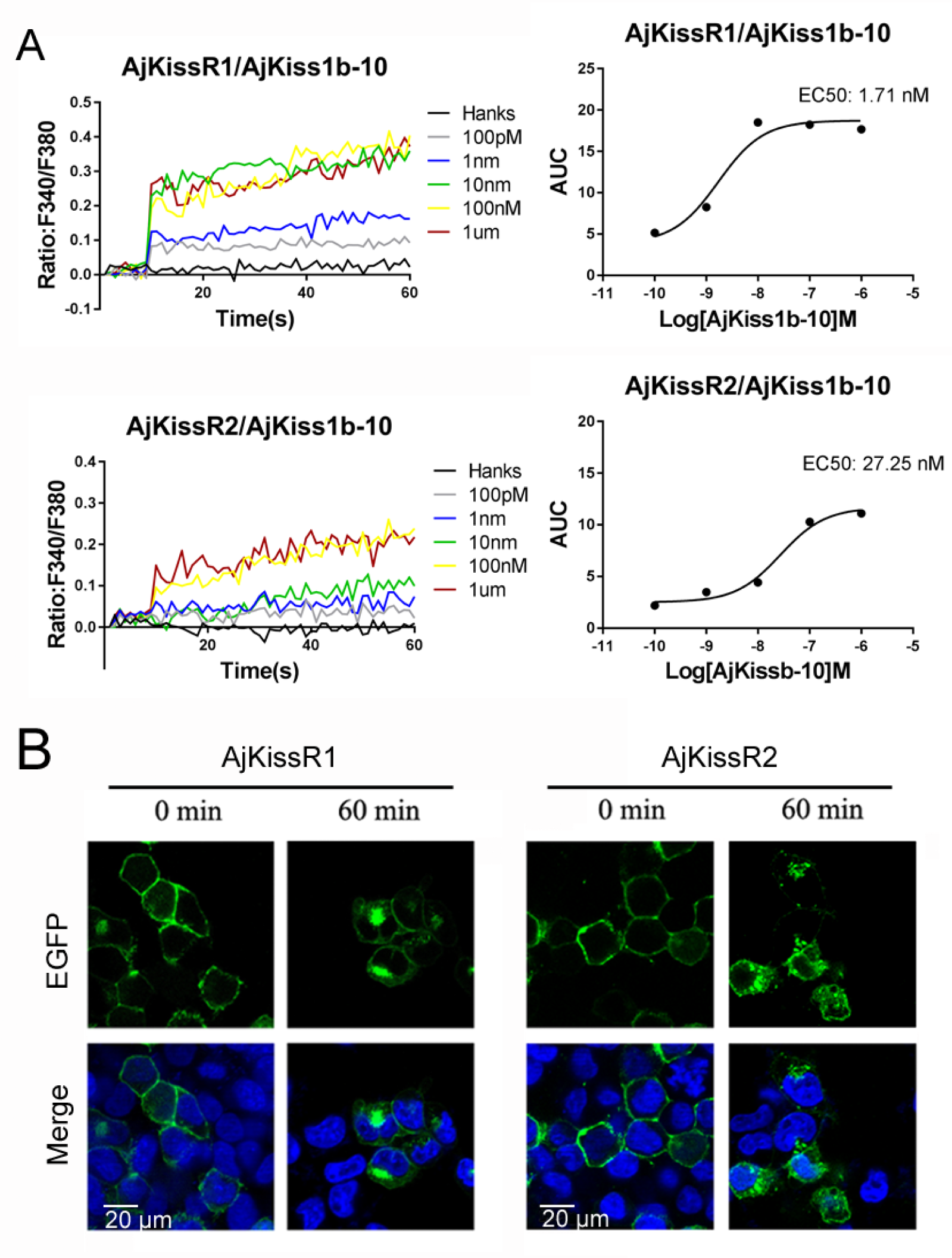
Functional activity of AjKiss1b-10. **A.** Intracellular Ca^2+^ mobilization in AjKissR1/2 expressing HEK293 cells was measured in response to AjKiss1b-10 with indicated concentrations using Fura-2/AM. **B.** Internalization of overexpressed AjKissR1/2 initiated by 1.0 μM AjKiss1b-10 in AjKissR1-EGFP or AjKissR2-EGFP expressing HEK293 cells was determined by confocal microscopy.

**Figure 6–figure supplement 1.**
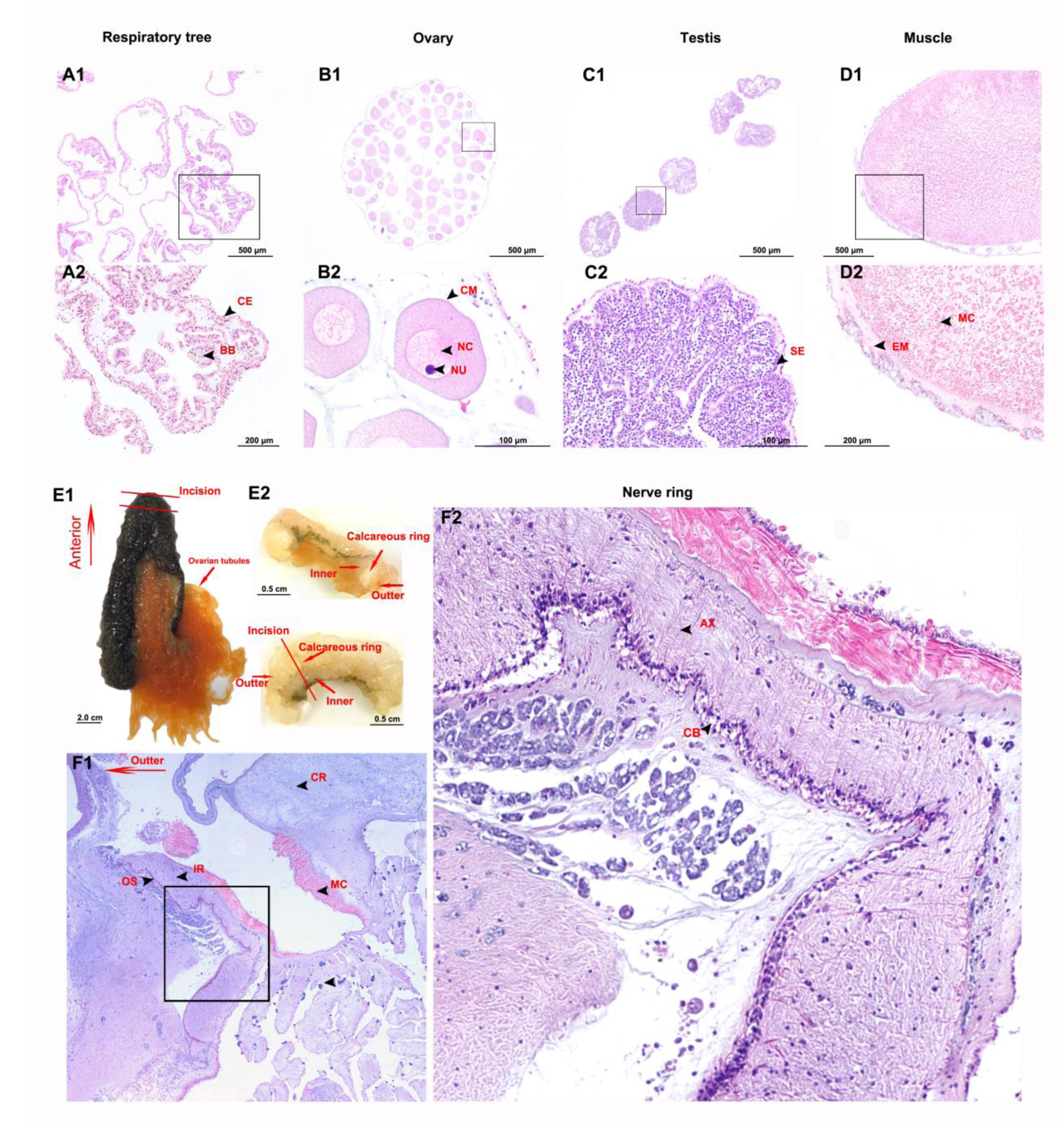
General morphology and histology of *Apostichopus japonicus* tissues. A-D. Light micrograhs of H&E staining section of respiratory tree, ovary, testis and muscle. (CE) coelomic epithelium, (BB) brown body, (CM) cell membrane, (NC) nucleus of oocytes, (NU) nucleolus of oocytes, (SE) spermatogenic epithelium, (EM) epithelium of muscle, (MC) myocyte. E. Gross anatomy of anterior part (ANP). F. Light micrograhs of H&E staining section of ANP and histology of nerve ring (NR). (OS) outter surface, (IR) internal region, (CR) calcareous ring, (AX) axon of neuron, (CB) cell body of neuron.

**Figure 6–figure supplement 2.**
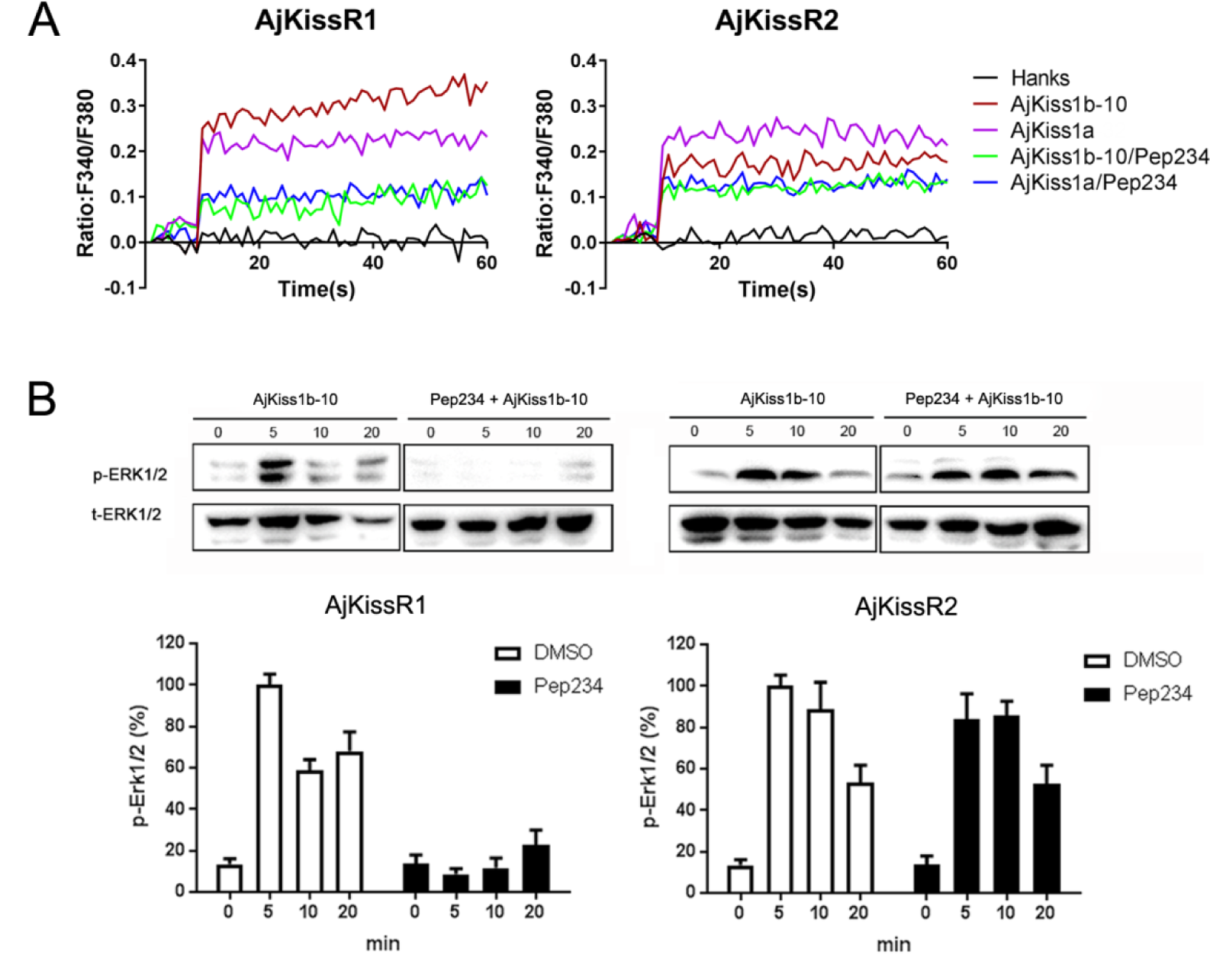
Inhibitory effect of pep234 on AjKissR1 and AjKissR2 activation. **A.** Intracellular Ca^2+^ mobilization in AjKissR1 and AjKissR2 expressing HEK293 cells was measured in response to 100 nM AjKiss1a or AjKiss1b-10 pretreated with DMSO or KISS1 antagonist pep234 (1 μM). **B.** ERK1/2 phosphorylation activity of Kps and inhibitory effect of pep234 AjKissR1 and AjKissR2 expressing HEK293 cells. Samples were measured after 2 h of ligand administration with or without pretreatment of pep234. Error bars represent SEM for three independent experiments.

**Figure 6–figure supplement 3.**
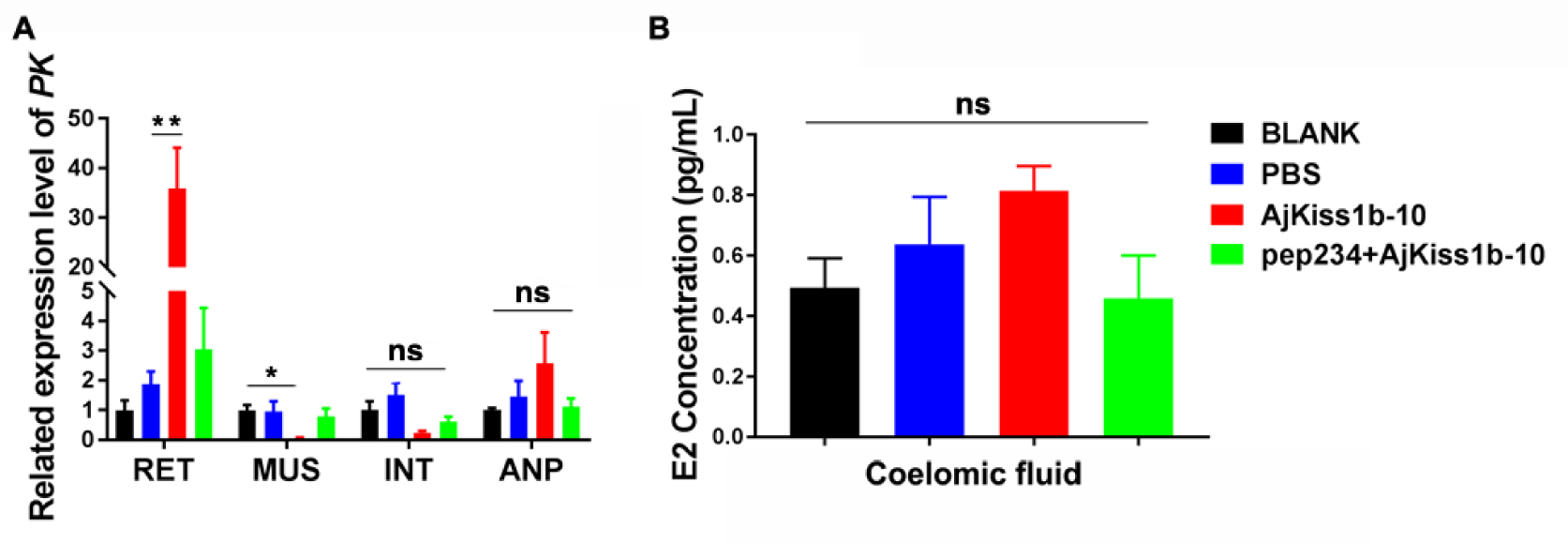
Functional activity of AjKiss1b-10 in *Apostichopus japonicus*. **A.** Gene expressional change of glycolytic enzyme gene *pyruvate kinase (PK)* in tissues of sea cucumbers responds to a 40-day administration of AjKiss1b-10. (RET) respiratory tree, (MUS) muscle, (INT) intestine, (ANP) anterior part of sea cucumber. **B.** E2 concentration in coelomic fluid of sea cucumbers did not significantly respond to AjKiss1b-10. Each symbol and vertical bar represents SEM (n=5). * indicates significant differences (P < 0.05), and ** indicates extremely significant differences (P < 0.01), ANOVA, Tukey‟s multiple comparison test.

**Figure 6–figure supplement 4.**
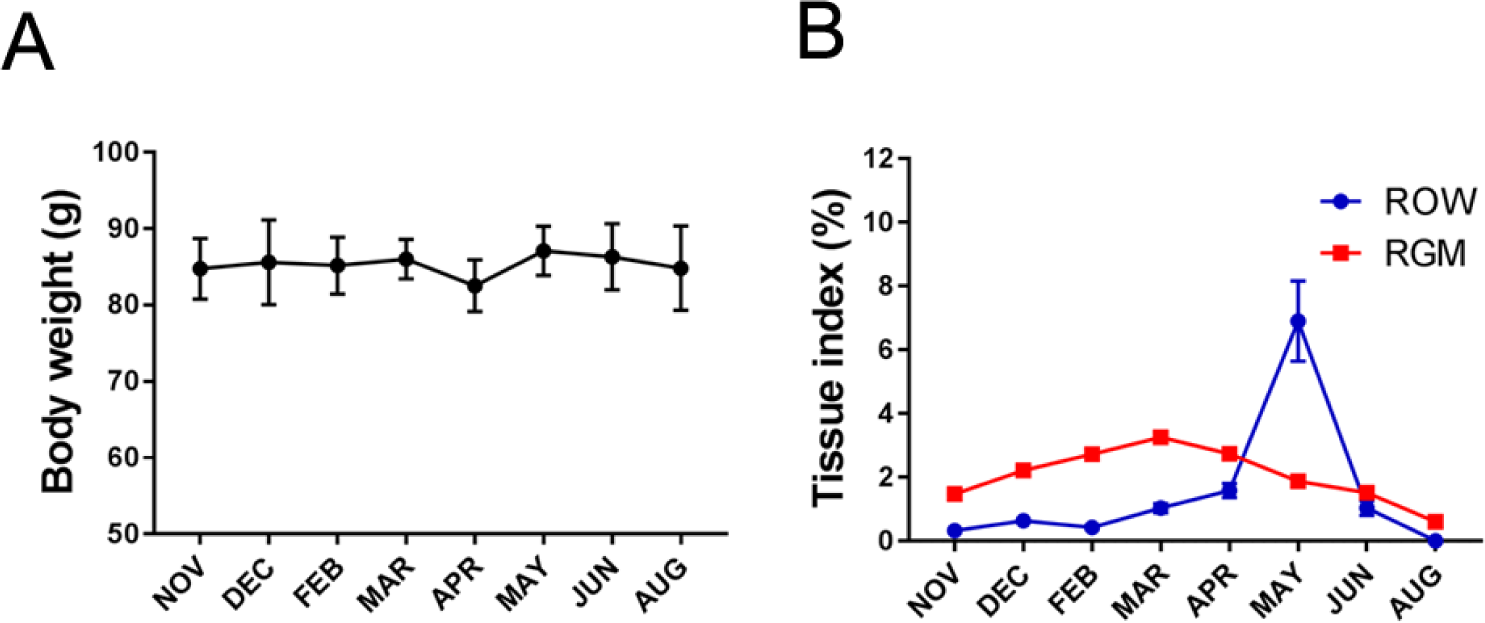
Mean body weight (**A**), relative gut mass (RGM), and relative ovary weight (ROW) (**B**) change over annual investigation. Each symbol and vertical bar represent SEM (n=5).

**Supplementary table 1.**
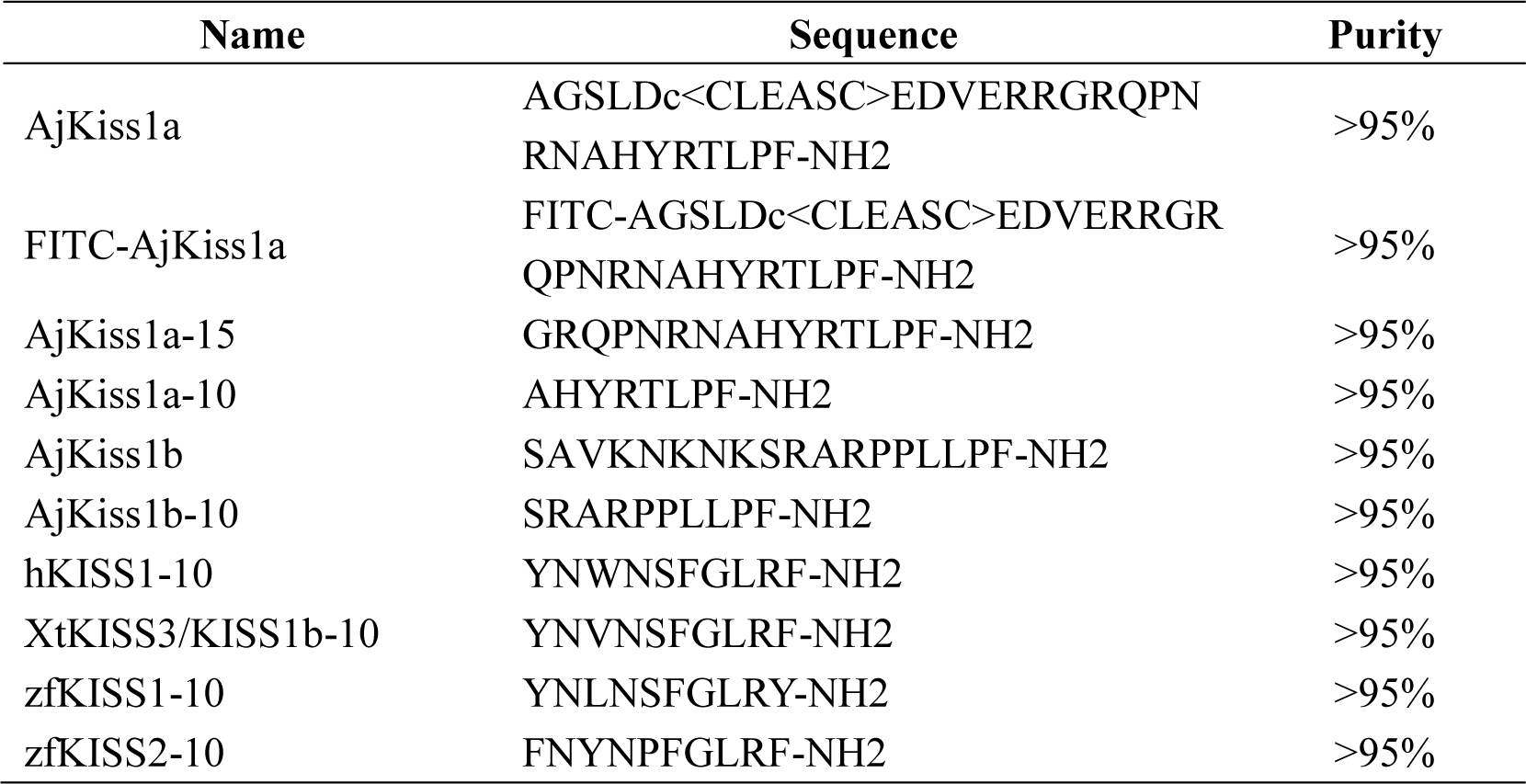
Sequence information of synthetic neuropeptide used. Note: c< > indicates disulfide bond

**Supplementary table 2.**
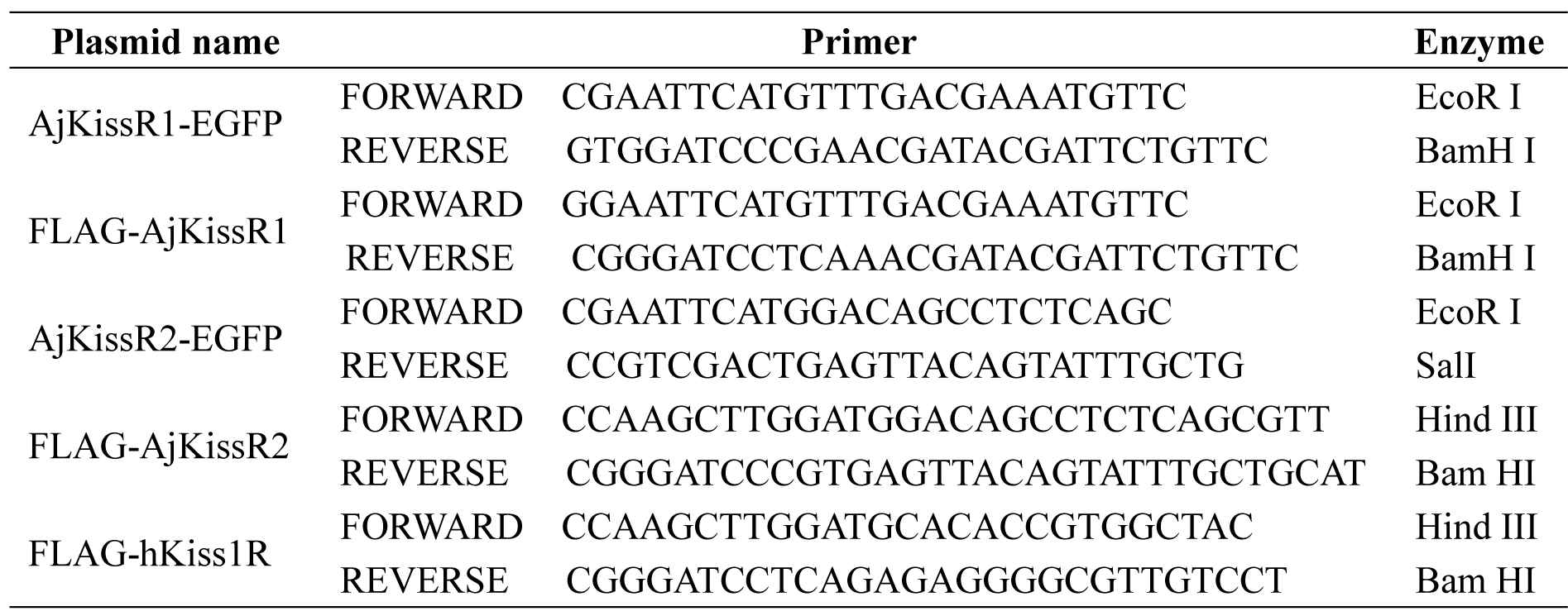
Primers for plasmid construction

**Supplementary Table 3.**
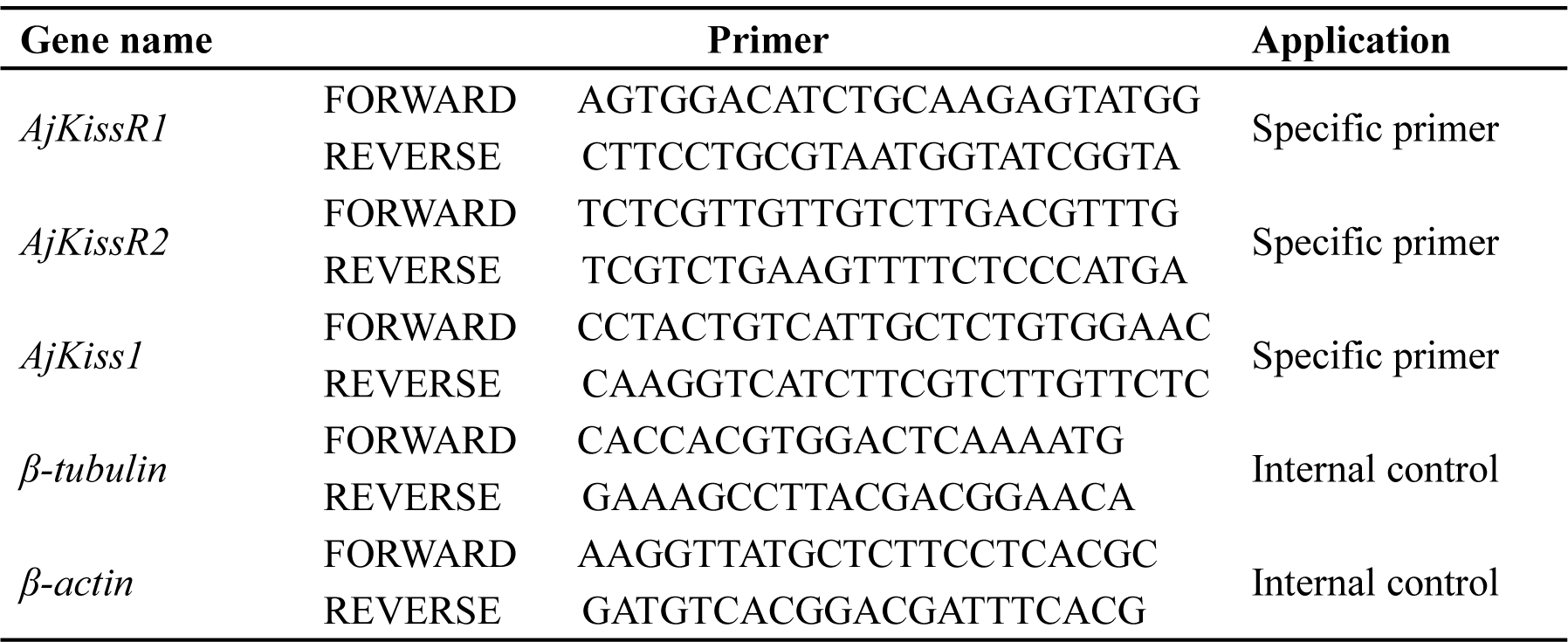
Primers for qPCR amplification

